# Force-activated zyxin assemblies coordinate actin nucleation and crosslinking to orchestrate stress fiber repair

**DOI:** 10.1101/2024.05.17.594765

**Authors:** Donovan Y.Z. Phua, Xiaoyu Sun, Gregory M. Alushin

## Abstract

As the cytoskeleton sustains cell and tissue forces, it incurs physical damage that must be repaired to maintain mechanical homeostasis. The LIM-domain protein zyxin detects force-induced ruptures in actin-myosin stress fibers, coordinating downstream repair factors to restore stress fiber integrity through unclear mechanisms. Here, we reconstitute stress fiber repair with purified proteins, uncovering detailed links between zyxin’s force-regulated binding interactions and cytoskeletal dynamics. In addition to binding individual tensed actin filaments (F-actin), zyxin’s LIM domains form force-dependent assemblies that bridge broken filament fragments. Zyxin assemblies engage repair factors through multi-valent interactions, coordinating nucleation of new F-actin by VASP and its crosslinking into aligned bundles by ɑ-actinin. Through these combined activities, stress fiber repair initiates within the cores of micron-scale damage sites in cells, explaining how these F-actin depleted regions are rapidly restored. Thus, zyxin’s force-dependent organization of actin repair machinery inherently operates at the network scale to maintain cytoskeletal integrity.

## INTRODUCTION

To functionally integrate into tissues, cells must perceive and respond to mechanical cues in their local environments (“mechanosense”). Mechanosensation is critical for development and tissue homeostasis^1^, and its dysfunction is associated with disease states such as hypertension^2^, fibrosis^3,4^, and cancer^5^. Cells physically engage their surroundings through cell-cell and cell-extracellular matrix adhesions that are linked to their intracellular actin cytoskeletons, notably contractile actin-myosin bundles known as stress fibers^6,7^. Stress fibers generate active forces that dynamically sculpt cell shape, facilitating cell migration^8,9^ and powering tissue rearrangements^10,11^. They also enable force transmission between the interior of cells and their tissue microenvironments, mediating the transduction of extracellular mechanical cues into intracellular biochemical signaling pathways (“mechanotransduction”)^12^. While dynamic regulation of stress fiber assembly, disassembly, and force generation capacity is required for morphogenesis^13^ and maintenance of tissue integrity^11^, feedback mechanisms which modulate the molecular composition and architecture of stress fibers as they fulfill their mechanical functions remain poorly understood.

Both exogenous^14–16^ and cell-intrinsic^17^ forces have the capacity to damage stress fibers, leading to F-actin depleted defects termed “stress fiber strain sites” (SFSSs)^17^. While complete SFSS rupture leads to loss of traction forces at a stress fiber’s associated adhesions^17^, the majority are repaired by zyxin, a force-responsive LIM (<u>L</u>IN-11, Isl-1, & <u>M</u>ec-3) domain protein critical for adhesion integrity^18–20^, durotaxis^21,22^, and development^23^. Live-cell imaging studies have shown zyxin initially localizes to SFSSs within seconds through its three C-terminal tandem LIM domains, producing micron-scale elongated puncta known as “flashes”^24^. Upon formation of a flash, zyxin’s N-terminal domain rapidly recruits the actin polymerization factor ENA/VASP (<u>Ena</u>bled/<u>Va</u>sodilator-<u>S</u>timulated <u>P</u>hosphoprotein, here termed ‘VASP’), followed by the actin cross-linking protein ɑ-actinin, mediating F-actin restoration and stress fiber repair on the timescale of minutes^17^. Recent work from our lab^25^ and others^26^ has shown that a subset of LIM domain proteins, including zyxin, directly bind individual tensed actin filaments through their tandem LIM domains. This suggests a model in which zyxin’s engagement of mechanically-strained F-actin is transduced into the activation of cytoskeletal repair machinery, a paradigmatic example of cytoskeletal mechanotransduction. While there is evidence for this sequence of events at the cellular level, the precise molecular mechanisms linking tensed F-actin recognition by zyxin’s LIM domains to the downstream effector functions of repair proteins remain unknown.

SFSS have been reported to be enriched with free plus (“barbed”) ends^17,27^ (where F-actin elongation preferentially occurs), presumed to be generated by filament breakage. This led to speculation that zyxin could coordinate F-actin elongation by VASP at these sites, as well as crosslinking of preexisting F-actin fragments by ɑ-actinin, to fill stress fiber gaps^17,27^. Such a “fill in” mechanism bears conceptual resemblance to damage repair in microtubules, where filament lattice gaps can be filled through the incorporation of free tubulin dimers^28,29^. However, features of the stress fiber repair system suggest this framework may be incomplete. First, both ɑ-actinin and VASP are oligomeric proteins featuring multiple F-actin binding domains, making it unclear how they would preferentially localize to F-actin remnants at SFSS versus adjacent F-actin-rich sites along the stress fiber. Second, SFSS can extend for several micrometers^17^, and it is not apparent how their F-actin depleted cores can sustain high forces without rupture for multiple minutes if restoration occurs primarily through elongation of peripheral broken filaments.

To probe the mechanism of zyxin-mediated stress fiber repair, here we pursued an *in vitro* reconstitution approach with purified proteins, culminating in a system which recapitulates restoration of ectopically damaged contractile F-actin bundles in the presence of active myosin motor forces. We find that zyxin forms force-dependent assemblies that bridge the remnants of mechanically ruptured actin filaments. These force-activated zyxin assemblies engage repair proteins through multivalent interactions, producing supramolecular networks that template *de novo* F-actin nucleation by VASP that is crosslinked into aligned bundles by ɑ-actinin. Super-resolution live-cell imaging studies suggest this network-level mechanism is the primary mode of stress fiber repair in cells, facilitating rapid restoration by initiating repair at the F-actin-depleted cores of SFSS, where they are most vulnerable to rupture.

## RESULTS

### Force-dependent zyxin assemblies span broken actin filament fragments

We recently reported that LIM domain proteins including zyxin bind in elongated puncta along single actin filaments in the presence of myosin motor forces, which we termed “patches”^25^. In total internal reflection fluorescence (TIRF) microscopy experiments, we noted that in a subset of patches, the intensity of the F-actin signal was dimmed relative to unbound filament regions. As the fluorophores and sparse F-actin labelling strategy we had employed rendered interpretation of this phenomenon ambiguous, here we updated our previously published actin force reconstitution assay^25^ (Figure 1A). We immobilized myosin Va (plus-end directed) and myosin VI (minus-end directed) on a passivated glass coverslip, followed by the addition of 100% ATTO-488 labelled F-actin and C-terminally Halo-tagged full-length zyxin labelled with Janelia Flour 646 (zyxin-Halo:JF646). This minimized the potential for fluorescence resonance energy transfer (FRET) between fluorophores and ensured that all F-actin in the system was labeled.

**Figure 1.**
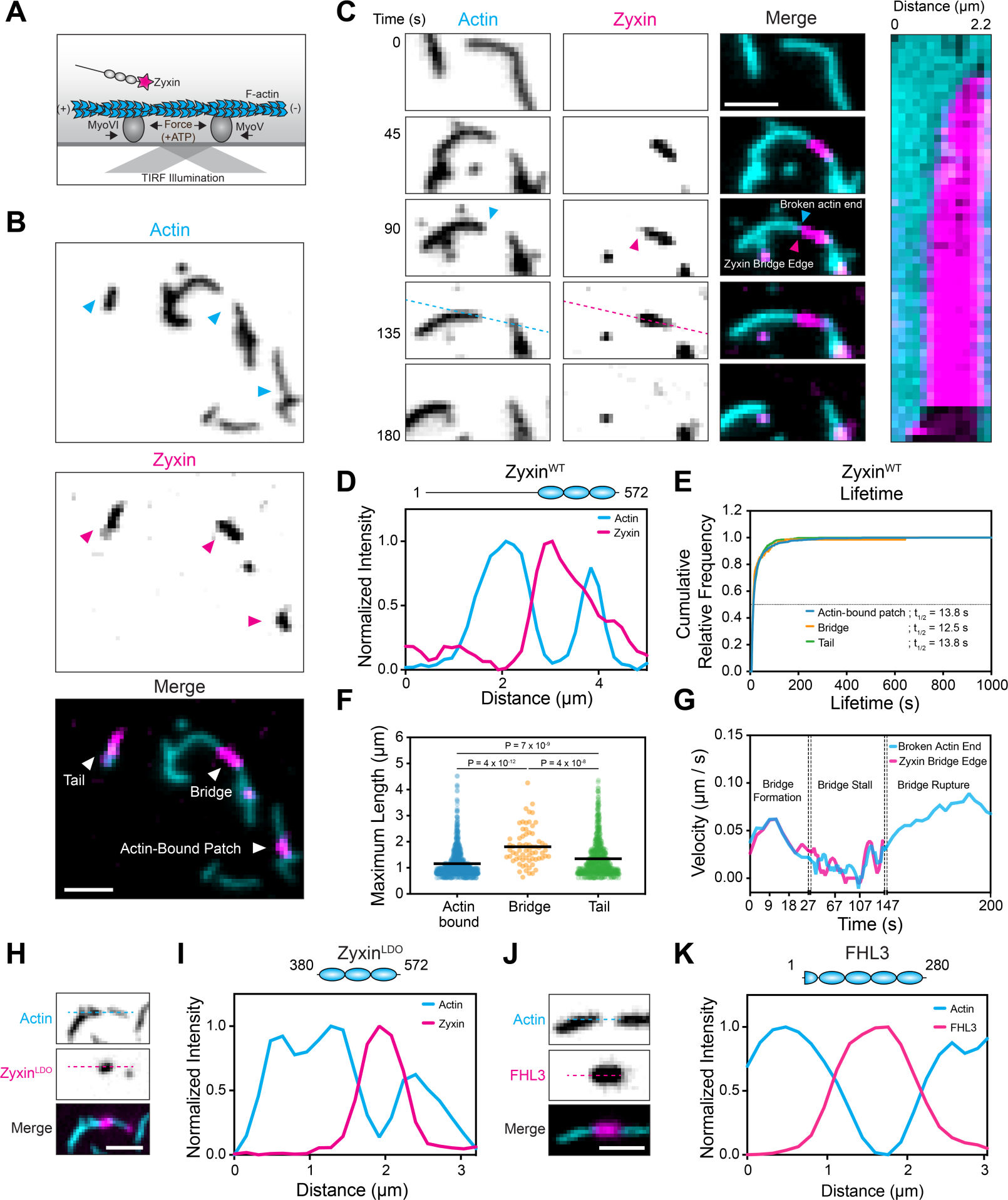
LIM protein bridges link fragments of broken actin filaments. **A)** Schematic of force reconstitution TIRF assay featuring fluorescently tagged zyxin. **B)** TIRF snapshot of 100% ATTO 488-labeled actin filaments (cyan) in the presence of 250 nM zyxin-Halo:JF646 (magenta) and myosin force generation. Arrowheads highlight different categories of zyxin patches. Scale bar, 2 μm. **C)** Montage (left) and kymograph (right) of zyxin bridge dynamics in the presence of 250 nM zyxin-Halo:JF646. Time labels correspond to both montage snapshots and kymograph. Scale bar, 2 μm. **D)** Normalized fluorescence intensity along dashed lines shown in C. **E)** Cumulative relative frequency of zyxin actin-bound patch, bridge, and tail lifetimes. Dotted line = 0.5. **F)** Maximum length of zyxin patches. Bars represent means. 68 ≤ n ≤ 925 from N = 3 biological replicates. Samples were compared by Turkey’s multiple comparisons test after ordinary one-way ANOVA. **G)** Instantaneous velocity over time of the interface between a zyxin bridge and a broken F-actin fragment, indicated with arrowheads in B. **H)** TIRF snapshot of a bridge formed in the presence of 250 nM zyxin^LDO^-Halo:JF646. Scale bar, 2 μm. **I)** Top: schematic of zyxin^LDO^ domain architecture. Bottom: normalized fluorescence intensity along dashed lines shown in H. **J)** TIRF snapshot of a bridge formed in the presence of 250 nM FHL3-Halo:JF646. Scale bar, 2 μm. **K)** Top: schematic of FHL3 domain architecture. Bottom: normalized fluorescence intensity along dashed lines shown in J.

Consistent with our previous findings^25^, in the presence of ATP, myosins apply stress along actin filaments, resulting in the formation of zyxin patches along strained actin filaments (here termed “F-actin-bound patches,” Figure 1B). However, we also observed a small fraction of zyxin patches that span between the ends of broken filament fragments in the absence of underlying F-actin, which we term “bridges” (Figures 1B-D, S1B-C). These bridges span an average length of 1.8 μm, with similar half-lives as actin-bound patches (Figures 1E-F). Based on the absence of F-actin signal and the lack of other soluble components in our system, we infer that bridges represent a force-dependent assembly of zyxin molecules. As zyxin bridges tether two filament fragments together, we hypothesized that they could act as molecular bandages, sustaining stress across broken filaments to prevent them from gliding apart due to motor activity. To test this hypothesis, we analyzed the dynamics of an F-actin fragment end and the zyxin bridge edge attached to it over time (Figure 1G). While the zyxin bridge is forming, the fragment end and the associated tip of the zyxin bridge glide at the same velocity. However, once the bridge reaches its maximum size, there is a decrease in velocity of both the zyxin bridge tip and the fragment end (a “bridge stall”). Eventually, the bridge rapidly dissociates in a single frame, accompanied by the fragment freely gliding away. Notably, the fragment end glides at a higher velocity after bridge rupture than prior to and during the bridge stall, suggesting that zyxin bridges are indeed force-sustaining structures that limit motor-driven F-actin translocation.

We also observed that zyxin bridges initially associated with two F-actin fragments can separate from one fragment while remaining associated with the other, forming structures we term “tails” (Figures 1B and S1D-E). Similar to bridges, filament motility stalls upon tails reaching their maximum size, which then accelerates upon tail dissociation (Figure S1F). This suggests tails are also under tension. It is unclear how tails adopt a load-bearing configuration in our assay, but we speculate that they represent zyxin patches which have become non-specifically associated with the glass coverslip. Regardless of the underlying mechanism, their existence provides additional support that zyxin can form force-dependent assemblies which are only partially associated with F-actin. As these end-associated structures grow in the same direction that their attached actin filament is gliding, we examined whether zyxin patches form in a directionally-dependent manner by immobilizing either myosin Va or myosin VI alone in our force reconstitution assay. In both conditions, we observed the formation of actin-bound patches, bridges, and tails. These data show zyxin patches form independently of F-actin polarity and motor directionality (Figure S1G), thereby providing a versatile molecular bandaging activity.

Previous work has demonstrated that zyxin’s isolated LIM domains are sufficient to form actin-bound patches^25,26^. To test whether they also support the formation of bridges, we examined a previously reported zyxin LIM-domain-only (zyxin^LDO^, amino acids 375-572) construct in our force reconstitution assay. We also observed the formation of bridges and tails with zyxin^LDO^ (Figures 1H-I) that featured similar properties to the wild-type (zyxin^WT^, Figures S1H-J), consistent with bridge formation being mediated by zyxin’s tandem LIM domains. As zyxin’s non-LIM sequence elements are dispensable, we hypothesized other LIM domains featuring conserved elements for force-activated actin binding could possess this activity^25^. Indeed, we find that C-terminally Halo-tagged four-and-a-half LIM domains 3 (FHL3) also forms bridges and tails with similar properties to both zyxin^WT^ and zyxin^LDO^ (Figures 1H-I and S1J-L). Collectively, these data suggest the formation of bridges is a conserved property of force-activated LIM domains.

### VASP is enriched on zyxin patches

We next examined how force-dependent zyxin assemblies interface with stress fiber repair machinery, first focusing on VASP. To test whether VASP directly associates with zyxin patches, we added both C-terminally Halo-tagged VASP labelled with Janelia Fluor 549 (VASP-Halo:JF549) and zyxin-Halo:JF646 to our force reconstitution assays (Figure 2A). In addition to actin-bound patches, we find that VASP robustly colocalizes with zyxin bridges and tails (Figures 2B-F). We observed two categories of binding events with distinct kinetics. Stable VASP binding events displayed near-perfect colocalization with zyxin along entire patches and feature lifetimes as long as their associated patch, whereas transient VASP binding events only partially cover zyxin patches and displayed flickering behavior with lifetimes shorter than 2 seconds (Figures 2G). These short-lived events feature binding dynamics reminiscent of those reported for VASP interacting with bare F-actin^30^.

**Figure 2.**
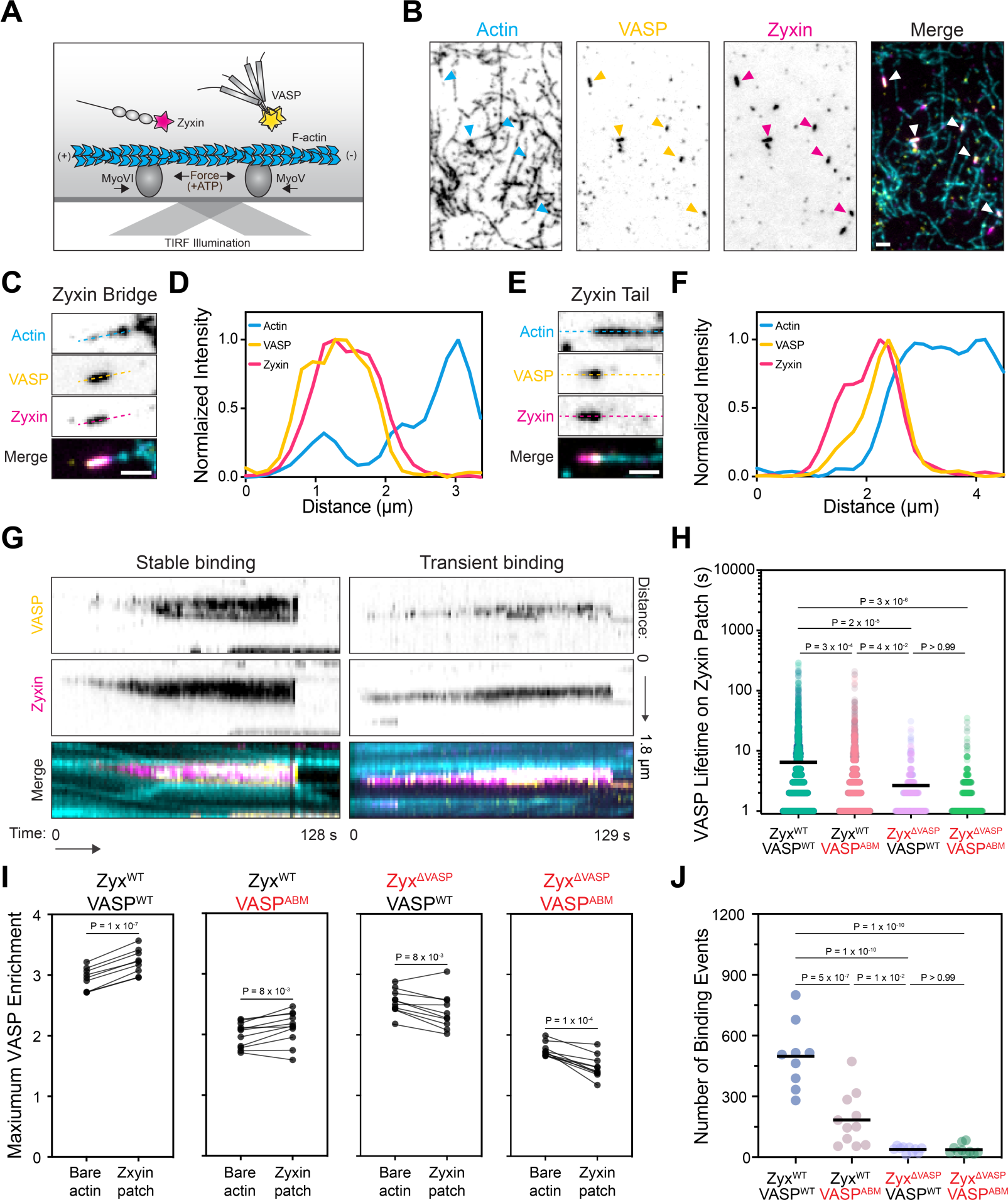
VASP is enriched on zyxin patches. **A)** Schematic of force reconstitution TIRF assay featuring fluorescently tagged zyxin and VASP. **B)** TIRF snapshot of VASP-enriched zyxin patches (arrowheads) formed in the presence of 30% ATTO 488-labeled actin filaments (cyan), 50 nM VASP-Halo:JF549 (yellow), and 25 nM zyxin-Halo:JF646 (magenta). Scale bar, 2 μm. **C)** Detail view of a zyxin bridge enriched with VASP. Scale bar, 2 μm. **D)** Normalized fluorescence intensity along dashed lines shown in C. **E)** Detail view of a zyxin tail enriched with VASP. Scale bar, 2 μm. **F)** Normalized fluorescence intensity along dashed lines shown in E. **G)** Kymographs highlighting two categories of VASP binding events on zyxin patches: stable binding (left) and transient binding (right). **H)** Lifetimes of VASP binding events on zyxin patches for indicated constructs. Bars represent means; 367 ≤ n ≤ 4479 from N = 2 biological replicates (represented by shades). Conditions were compared by Turkey’s multiple comparisons test after ordinary one-way ANOVA. **I)** Maximum VASP enrichment on zyxin patches versus bare F-actin during equal imaging periods across trials for indicated constructs. 9 ≤ n ≤ 11 from N = 2 biological replicates, compared by paired t test. **J)** Number of VASP-zyxin binding events detected during equal imaging periods across trials for wild-type and mutant constructs. 9 ≤ n ≤ 11 from N = 2 biological replicates. Conditions were compared by Turkey’s multiple comparisons test after ordinary one-way ANOVA.

We furthermore observed that VASP preferentially localized to the F-actin-associated ends of tails (Figure 2E,F), suggesting both zyxin binding and F-actin binding could contribute to VASP’s zxyin patch engagement. To dissected the interplay of these activities, we examined a VASP mutant deficient in both G- and F-actin binding^30^ [VASP^ABM^: L226A, I231A, L235A, R(273-275)E, K276E], as well as a zyxin mutant deficient in recruiting VASP to SFSS in cells^17,31^ [zyxin^ΔVASP^: F(75, 97, 108, 118)A] (Figure S2A). The half-lives of zyxin patches were highly similar across all conditions we examined with these constructs, suggesting that VASP recruitment does not detectably impact patch stability in our assay (Figure S2B). However, VASP^ABM^ binding events displayed a significantly shorter mean lifetime on zyxin^WT^ patches than VASP^WT^ binding events, an effect which was enhanced in all conditions featuring zyxin^ΔVASP^ (Figure 2I). In the zyxin^WT^ / VASP^WT^ condition, we observed a substantial enrichment of VASP on zyxin patches versus bare F-actin regions. This VASP enrichment persisted but was significantly diminished in the zyxin^WT^ / VASP^ABM^ condition. In both the zyxin^WT^ / VASP^WT^ and zyxin^WT^ / VASP^ABM^ conditions, we observed many long-lived VASP binding events on patches. However, in all conditions featuring zyxin^ΔVASP^, VASP enrichment on zyxin patches was essentially eliminated, and only transient binding events were observed (Figures 2I and S2A, C-D). This suggests that long-lived VASP association with patches is primarily mediated by zyxin-VASP binding interactions. Similar trends were observed when we quantified the number of VASP binding events on zyxin patches (Figure 2J). Together, these data suggest zyxin-VASP binding is the primary interaction mediating VASP recruitment to zyxin patches, while VASP-F-actin binding plays a reinforcing role. Collectively, our results show that force-dependent zyxin patches can directly concentrate VASP.

### Zyxin engages ɑ-actinin pre-bound to F-actin, which can mediate filament zippering

To probe the interaction between zyxin patches and ɑ-actinin, we next studied the interplay between C-terminally Halo-tagged ɑ-actinin-1 labelled with Janelia Flour 549 (ɑ-actinin-Halo:JF549) and zyxin-Halo:JF646 in force reconstitution assays (Figure 3A). Despite some degree of co-localization (Figure 3B), unlike VASP (Figure 2B-D), we did not observe extensive recruitment of ɑ-actinin to zyxin patches. We reasoned the partial overlap of some ɑ-actinin puncta with zyxin patches could be mediated either by indirect colocalization of zyxin patches and ɑ-actinin to the same region on F-actin or by a weak direct interaction between the proteins. To dissect these possibilities, we pursued a parallel approach to our VASP studies, examining an ɑ-actinin F-actin binding mutant^32^ (ɑ-actinin^ABM^: I110A, I115A) and a zyxin mutant featuring deficient ɑ-actinin recruitment to SFSS in cells^17^ (zyxin^Δɑ-actinin^: Δ1-42, Figures S2A-B). We observed no significant difference in the number of colocalized events between the zyxin^WT^ / ɑ-actinin^WT^ and zyxin^Δɑ-actinin^ / ɑ-actinin^WT^ conditions, suggesting zyxin binding does not drive recruitment of soluble ɑ-actinin to patches (Figure 2E). Consistently, there was negligible colocalization of ɑ-actinin^ABM^ with either zyxin^WT^ or zyxin^Δɑ-actinin^, suggesting ɑ-actinin’s actin-binding activity is required for patch association. However, when focusing on zyxin–ɑ-actinin co-localization events, we did observe modest but significantly increased enrichment of ɑ-actinin^WT^ on patches versus bare F-actin when comparing the zyxin^WT^ and zyxin^Δɑ-actinin^ conditions. This suggests that zyxin:ɑ-actinin binding is indeed occurring, likely by reinforcing patch association of F-actin-bound ɑ-actinin rather than by initiating recruitment.

**Figure 3.**
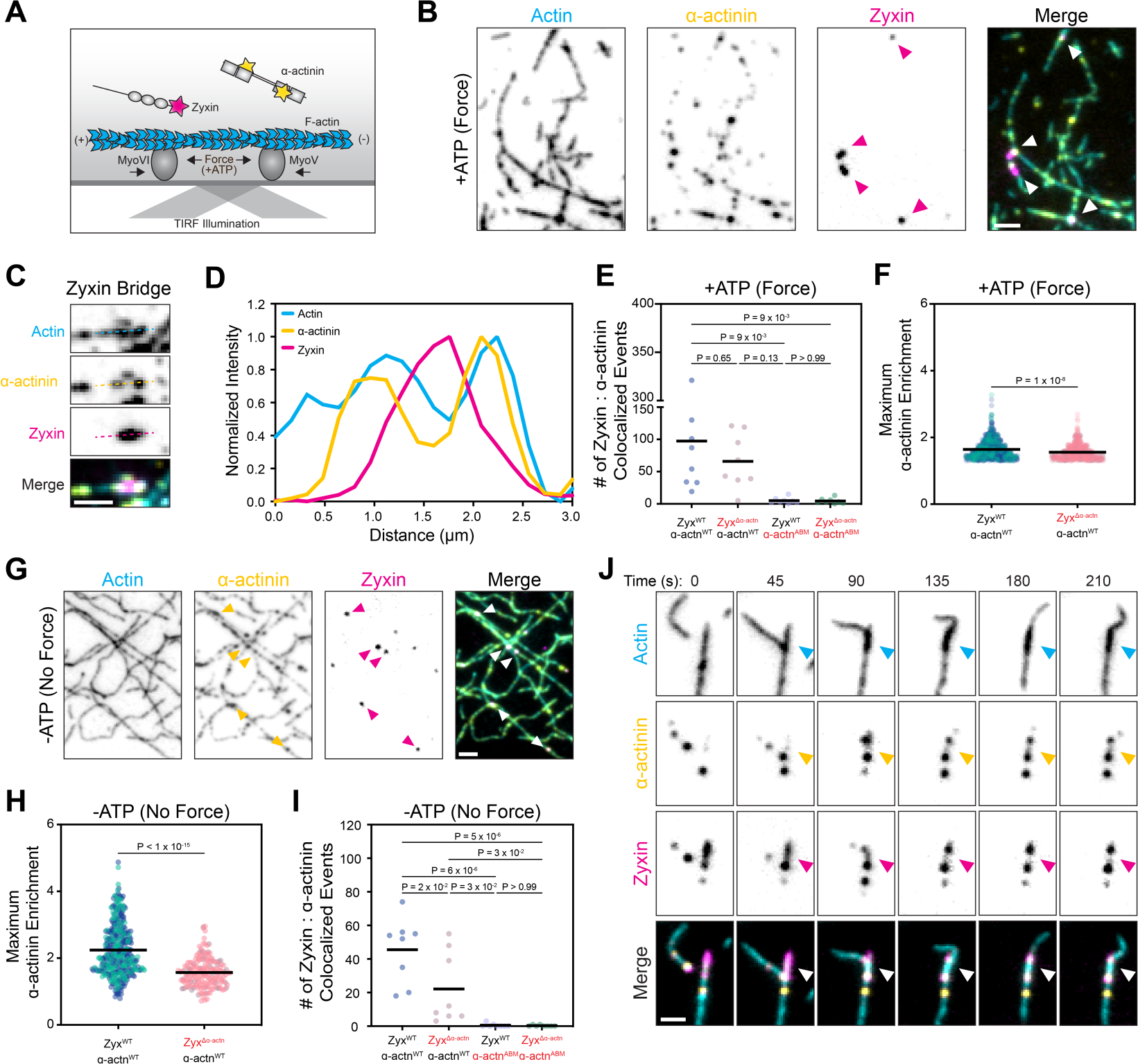
ɑ-actinin mediates actin filament bundling along zyxin patches. **A)** Schematic of force reconstitution TIRF assay featuring fluorescently tagged zyxin and ɑ-actinin. **B)** TIRF snapshot of ɑ-actinin clusters co-localized with zyxin patches and clusters (arrowheads) in the presence of immobilized myosin forces (+ATP). Assay was performed with 30% ATTO 488-labeled actin filaments (cyan), 50 nM ɑ-actinin-Halo:JF549 (yellow), and 25 nM zyxin-Halo:JF646 (magenta). Scale bar, 2 μm. **C)** Detail view of a zyxin patch featuring associated ɑ-actinin clusters. Scale bar, 2 μm. **D)** Normalized fluorescence intensity along dashed lines shown in C. **E)** Number of ɑ-actinin–zyxin colocalization events detected during equal imaging periods across trials for the indicated constructs in the presence of immobilized myosin forces (ATP); n = 8 from N = 2 biological replicates. Conditions were compared by Turkey’s multiple comparisons test after ordinary one-way ANOVA. **F)** Maximum wild-type ɑ-actinin enrichment per zyxin^WT^ (n = 779) or zyxin^Δɑ-actinin^ (n = 527) patch colocalization event in the presence of immobilized myosin forces (+ATP). Bars represent means; N = 2 biological replicates (represented by shades). Conditions were compared by unpaired t test. **G)** TIRF snapshot of zyxin clusters colocalized with ɑ-actinin. Assay was performed as in B, but in the absence of force generation (-ATP). Scale bar, 2 μm. **H)** Number of ɑ-actinin-zyxin colocalization events detected during equal imaging periods across trials for the indicated constructs in the absence of immobilized myosin forces (-ATP). n = 8 from N = 2 biological replicates (represented by shades). Conditions were compared by Turkey’s multiple comparisons test after ordinary one-way ANOVA. **I)** Maximum wild-type ɑ-actinin enrichment per zyxin^WT^ (n = 364) or zyxin^Δɑ-actinin^ (n = 177) cluster colocalization event in the absence of immobilized myosin forces (-ATP). Bars represent means; N = 2 biological replicates (represented by shades). Conditions were compared by unpaired t test. **J)** Montage of a gliding actin filament (cyan) zippering along a zyxin patch (magenta) through ɑ-actinin clusters (yellow). Arrowheads highlight the position of the zyxin patch. Assay was performed with 30% ATTO 488-labeled actin filaments, 50 nM ɑ-actinin-Halo:JF549, and 250 nM zyxin-Halo:JF646. Scale bar, 2 μm.

In addition to ɑ-actinin’s recruitment to SFSS through zyxin, the two proteins co-localize at diffraction-limited puncta throughout stress fibers^17^. We therefore hypothesized that zyxin could reciprocally be recruited to F-actin through ɑ-actinin’s force-independent F-actin binding activity. To test this hypothesis, we examined zyxin^WT^ and ɑ-actinin^WT^ in our assay in the absence of ATP to preclude the formation of force-dependent zyxin patches. We observed the co-localization of zyxin puncta with ɑ-actinin clusters on F-actin, supporting a force-independent association (Figures 3G and S3B). Zyxin^Δɑ-actinin^ displayed significantly reduced colocalization with ɑ-actinin^WT^ under these conditions by multiple metrics (Figure 2H,I), suggesting this interaction is mediated by direct binding between the proteins. As anticipated, negligible F-actin-associated zyxin puncta were observed in the presence of ɑ-actinin^ABM^ (Figure 2H), confirming force-independent F-actin colocalization of the proteins is completely dependent on ɑ-actinin’s F-actin binding activity.

The co-existence of force-dependent and force-independent zyxin–ɑ-actinin interaction modes on F-actin, as well as ɑ-actinin’s capacity to crosslink filaments, led us to examine the interplay between these activities under force-generating (+ATP) conditions. With zyxin^WT^ and ɑ-actinin^WT^, we observed the contemporaneous formation of zyxin patches, F-actin gliding, and F-actin bundling. Strikingly, we found that ɑ-actinin puncta on one actin filament mediated lateral association with a zyxin patch on another filament when myosin-driven gliding brings them into contact (Figure 3J and S3D), leading to co-linear ‘zippering’ of the two filaments along the patch. This activity could facilitate mechanical reinforcement of SFSS by locally coupling damaged filaments to undamaged ones, consistent with prior speculation^17^, while maintaining the appropriate inter-filament geometry of the stress fiber. Overall, these data show that zyxin can interact with F-actin-bound ɑ-actinin to reinforce damage sites through F-actin bundling.

### Zyxin, VASP, and ɑ-actinin associate through multivalent interactions

We next investigated the mechanistic basis of zyxin’s interactions with VASP and ɑ-actinin. VASP is a homotetramer, ɑ-actinin is a homodimer, and zyxin’s N-terminus features four VASP-binding ActA repeats, highlighting the potential for multivalent interactions in this system. Previous studies have shown that VASP has the capacity to form droplets which facilitate F-actin polymerization^33^, while the zyxin homolog LIMD1 partitions into micron-scale puncta in cells when ectopically induced to dimerize with an optogenetic system^34^. Upon ɑ-actinin-mediated bundling of two filaments at a patch site, we noted that the zyxin within the patch redistributed into two ɑ-actinin clusters (Figure 3J), reminiscent of de-wetting phenomena reported in microtubule-associated clusters of augmin^35,36^. We therefore examined whether zyxin, VASP, and ɑ-actinin have the capacity to form droplets *in vitro*, a hallmark of multivalent interaction-driven clustering^37^.

We first mixed zyxin, VASP, and ɑ-actinin individually with 3% PEG 8K and monitored droplet formation. At a concentration of 1 μM, only VASP forms droplets in isolation (Figure S4A), consistent with a previous report^33^. However, combining the proteins together resulted in the concentration-dependent formation of micron-scale droplets featuring all three components (Figures S4B-C). We observe fusion events between spontaneously contacting droplets, as well as signal recovery of all three proteins in fluorescence recovery after photobleaching (FRAP) assays, suggesting zyxin–VASP–ɑ-actinin droplets feature dynamics similar to other multivalent supramolecular assemblies (Figures S4D-F)^38^. We also observed small clusters at native cellular concentrations of the proteins^39^ in the absence of PEG 8K (Figure S4G), suggesting multivalent clustering can occur under physiological conditions.

We next dissected the molecular interactions mediating droplet formation. To assess which sequence modules of zyxin were necessary, we mixed VASP and ɑ-actinin with a minimal construct containing zyxin’s N-terminal binding motifs for these factors (amino acids 1-160, zyxin^N-term^) or with zyxin^LDO^ (Figures S4H-I). Consistent with its established role in mediating interactions with actin regulatory machinery in cells^17,25^, zyxin^N-term^ was sufficient to promote droplet formation, while zyxin^LDO^ did not localize to droplets. We next probed pairwise interactions between the proteins. At 1 μM each of zyxin and VASP, we observed the formation of bipartite droplets (Figure S4J-K). Versus zyxin^WT^, zyxin^N-term^ displayed modestly diminished enrichment in droplets, while zyxin^ΔVASP^’s enrichment was strongly decreased. This suggests that zyxin’s enrichment in droplets is primarily mediated through direct binding to VASP. As anticipated from the three-protein experiments, zyxin^LDO^ did not partition into droplets with VASP. Unlike the robust VASP droplet formation observed under all conditions we tested, zyxin and ɑ-actinin formed bipartite droplets only in the presence of zyxin^WT^, with no droplets observed in conditions featuring zyxin^Δɑ-actinin^, zyxin^N-term^, or zyxin^LDO^ (Figure S5L-M). This suggests intact zyxin is required, either due to the presence of additional ɑ-actinin binding sites beyond the N-terminal domain, or to mediate stabilization of the N-terminal domain’s interactions with ɑ-actinin. ɑ-actinin^ABM^ does support droplet formation with zyxin^WT^ at high (1 μM) protein concentrations in the presence of crowding agents. This suggests ɑ-actinin’s actin-binding activity likely promotes the formation of bipartite ɑ-actinin–zyxin clusters on F-actin, either by locally concentrating ɑ-actinin (we employed 25-50 nM in our force reconstitution assays), or by promoting the cytoskeletal localization of clusters, rather than by directly gating their formation. Taken together, these data suggest that multivalent interactions between specific binding motifs mediate zyxin’s formation of supramolecular assemblies with actin regulatory proteins.

### VASP-enriched zyxin patches drive actin nucleation and polymerization

To probe how zyxin’s interactions with actin regulatory machinery could contribute to F-actin restoration at SFFSs, we next examined their effects on actin polymerization *in vitro*. VASP droplets were recently reported to concentrate G-actin and capture assembling actin filaments^33^, suggesting VASP associated with zyxin patches might drive local F-actin polymerization. We therefore incorporated profilin-G-actin (ATTO-488 labeled) into our force reconstitution assay alongside VASP-Halo:JF549 and zyxin-Halo:JF646 (Figure 4A) and monitored actin dynamics. To provide favorable conditions for filament elongation at plus-ends marked by zyxin patches, we initially examined preparations solely featuring myosin Va.

**Figure 4.**
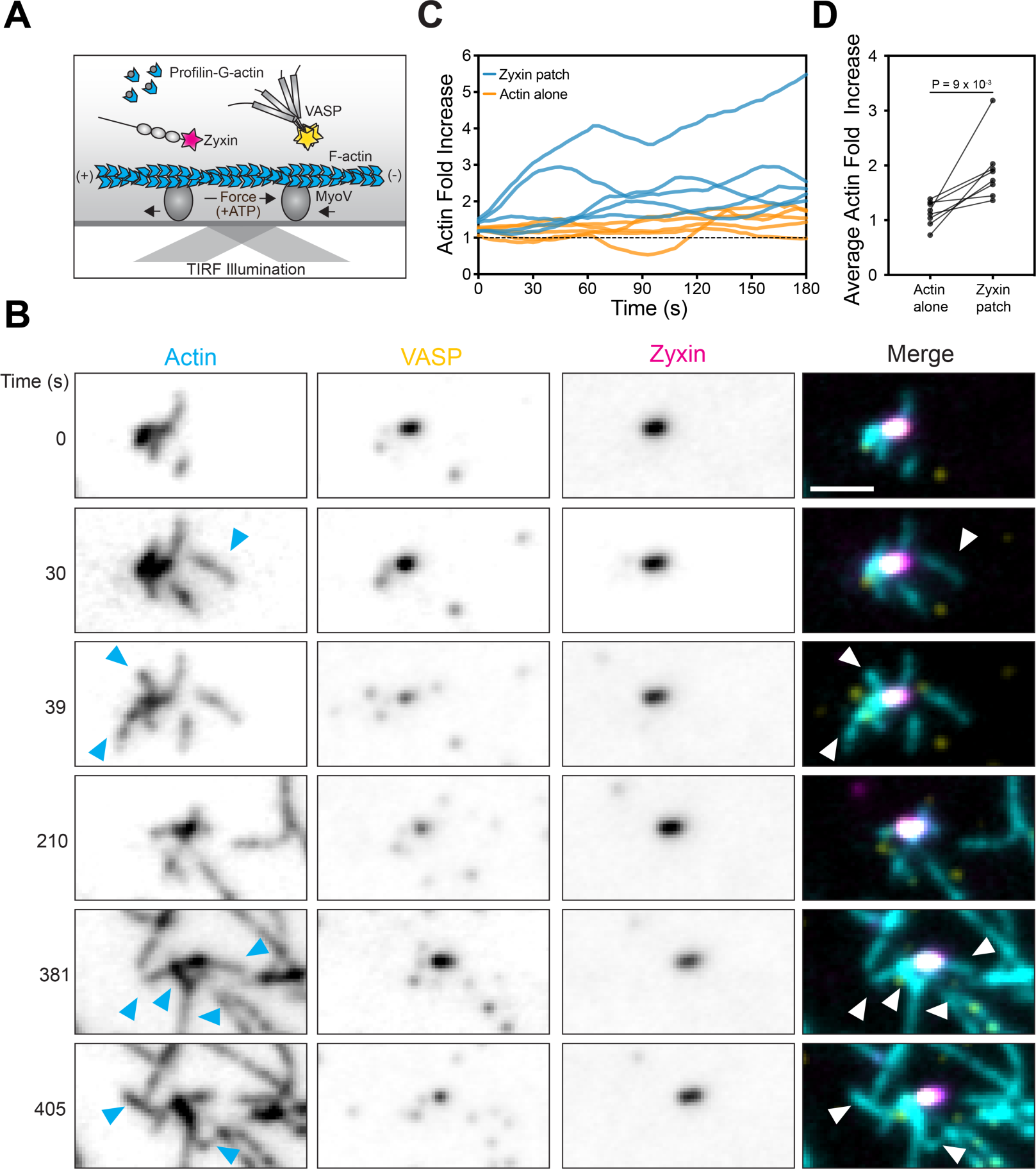
VASP nucleates F-actin when engaged at zyxin patches. **A)** Schematic of force reconstitution TIRF assay featuring fluorescent zyxin, VASP, and profilin–G-actin. **B)** Montage of actin filaments (cyan, 30% ATTO 488-labeled) being nucleated and elongating out of a VASP-enriched (yellow) zyxin patch (magenta) while being extruded by immobilized myosin V forces (+ATP). Arrowheads highlight examples of actin nucleation / polymerization. Assay was performed with 0.5 μM 30% ATTO-488 G-actin, 2 μM profilin, 50 nM VASP-Halo:JF549, and 25 nM zyxin-Halo:JF646. Scale bar, 2 μm. **C)** Quantification of actin intensity fold increase over time at VASP-enriched zyxin patches versus bare F-actin regions. Assay conditions were as in B. Dashed line = 1. **D)** Pairwise comparison of the average actin intensity fold increase at VASP-enriched zyxin patches vs. bare F-actin regions across trials using assay conditions as in B. n = 8 trials (examining 1-6 fields of view per trial) from N = 2 biological replicates, compared by paired t test.

Strikingly, while some elongation of pre-existing filaments did occur at VASP-enriched zyxin patches, we instead primarily observed extensive nucleation of new filaments (Figures 4A,B). These filaments were extruded by motor forces and fragmented as they polymerized, with no apparent preferential orientation relative to the source patch. This suggests VASP’s F-actin-binding activity is insufficient to bundle filaments or resist motor-driven sliding at zyxin patches. Quantification showed a significantly greater increase in actin fluorescence signal over time in the vicinity of VASP-enriched patches versus F-actin devoid of patches, suggesting concentration of VASP by zyxin patches promotes F-actin assembly (Figures 4C-D and S5A). Preparations featuring myosin VI instead of myosin Va featured highly similar F-actin polymerization dynamics, suggesting patches generated by minus-end directed forces can similarly promote VASP-mediated nucleation (Figures S5B-D). Together, these data show zyxin patches serve as a force-dependent platform for promoting F-actin assembly by VASP, primarily through *de novo* filament nucleation.

### Zyxin-associated machinery is sufficient to build and repair contractile networks

Our studies of zyxin’s interactions with cytoskeletal partners suggest that patches on individual filaments can serve as hubs to focus key activities regulating cytoskeletal dynamics. As stress fibers are multi-filament assemblies, we next examined how these activities are coordinated at the cytoskeletal network scale to execute stress fiber repair. We first analyzed a mixture of zyxin-Halo:JF646, VASP-Halo:JF549, ɑ-actinin-Halo (unlabeled), and profilin-G-actin (ATTO-488 labeled) in a force reconstitution assay featuring immobilized myosin VI and pre-polymerized F-actin (Figure 5A). Over the course of ten minutes, gliding of actin filaments stalled, followed by their apparent thickening to form nonmotile F-actin bundles which remained anchored within the TIRF field (Figures 5B, S6A-B).

**Figure 5.**
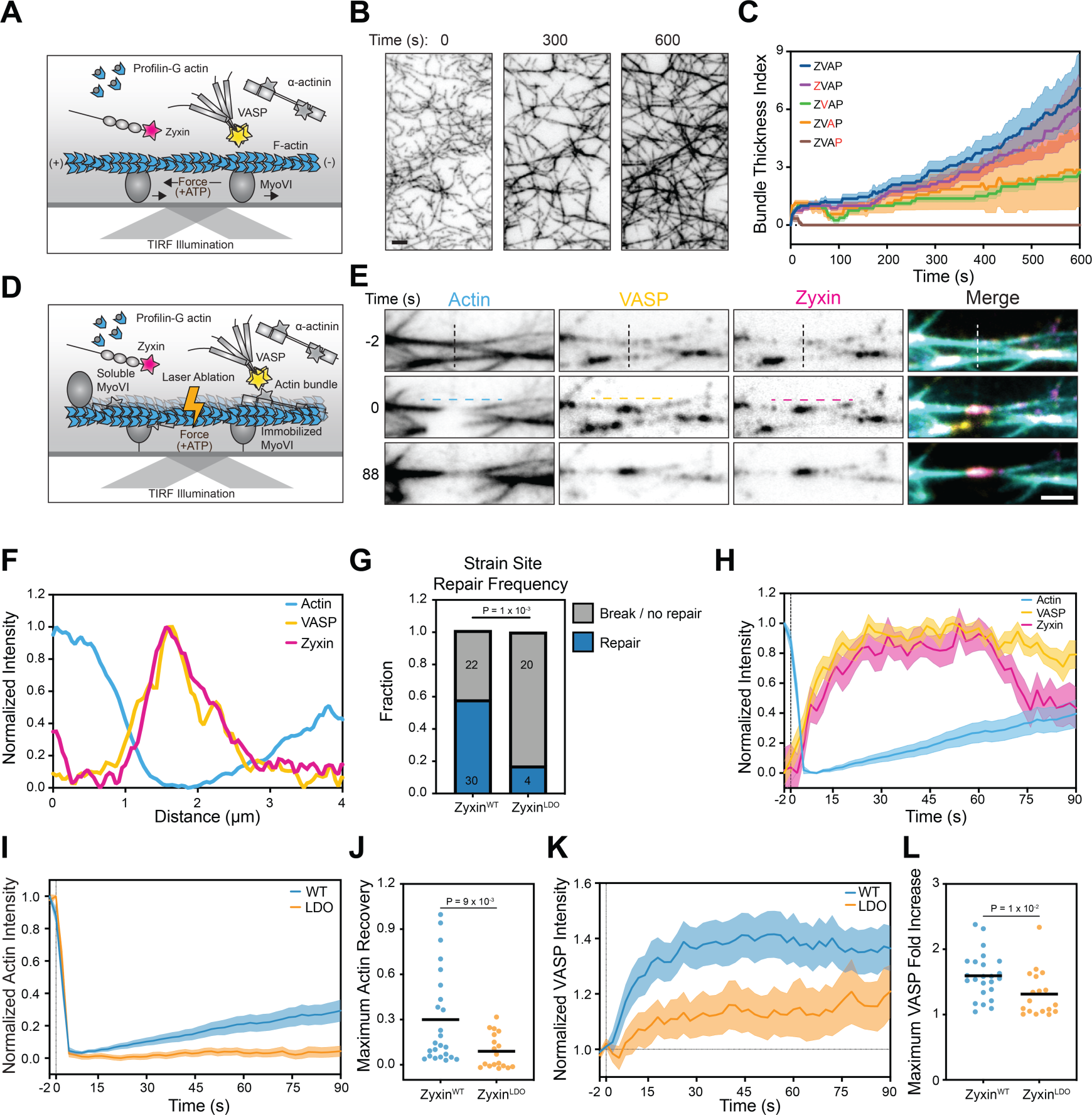
Reconstitution of mechanically damaged stress fiber repair. **A)** Schematic of bundle reconstitution TIRF assay featuring fluorescent zyxin, VASP, and profilin-G-actin, as well as unlabeled ɑ-actinin. **B)** Montage of F-actin bundles (ATTO-488 signal) thickening over time. The assay was performed with 25 nM zyxin-Halo:JF646, 50 nM VASP-Halo:JF549, 50 nM ɑ-actinin-Halo, 0.5 μM 30% ATTO-488 labeled G-actin, and 2 μM profilin. Scale bar, 2 μm. **C)** Quantification of bundle thickness index versus time in assays performed as in B, where red colored letters denote experiments featuring the absence of the indicated proteins (‘Z’, zyxin; ‘V’, VASP; ‘A’, ɑ-actinin; ‘P’, profilin-G-actin). Data are presented as mean ± SEM; 7 ≤ n ≤ 11 trials from N = 2 biological replicates. **D)** Schematic of contractile actin-myosin bundle reconstitution via the inclusion of soluble myosin VI, facilitating photoablation TIRF assays. Experiments feature fluorescent zyxin, VASP, and profilin-G-actin, as well as unlabeled ɑ-actinin. **E)** Montage of laser ablation of a contractile bundle, triggering a zyxin flash and subsequent F-actin recovery. Vertical dashed line indicates region targeted for ablation at time = 0 s. Assay was performed with 100 nM zyxin-Halo:JF646, 50 nM VASP-Halo:JF549, 100 nM ɑ-actinin-Halo, 1 μM 30% ATTO-488 labeled G-actin, and 4 μM profilin. Scale bar, 2 μm. **F)** Normalized fluorescence intensity line scans along the zyxin flash immediately below the horizontal dashed lines shown in E. **G)** Strain site repair frequency in the presence of zyxin^WT^ versus zyxin^LDO^. Number of strain sites in each category are indicated; N ≥ 3 biological replicates. Conditions were compared by Fisher’s exact test. **H)** Normalized fluorescence intensity over time of indicated proteins at strain sites that underwent repair in the presence of zyxin^WT^. Dashed line indicates time = 0 seconds, when laser ablation occurred. Data are presented as mean ± SEM. **I-L)** Analysis of pooled strain sites (repaired and not repaired) observed in the presence of zyxin^WT^ (n = 24 from N = 5 biological replicates) or zyxin^LDO^ (n = 18 from N = 4 biological replicates). **I)** Normalized actin fluorescence intensity over time. Vertical dashed line indicates time = 0 seconds, when laser ablation occurred. Data are presented as mean ± SEM. **J)** Comparison of maximum actin recovery as fraction of initial intensity (unpaired t test). **K)** Normalized VASP fluorescence intensity fold increase over time. Vertical dashed line indicates time = 0 seconds, when laser ablation occurred. Horizontal dashed indicates baseline VASP intensity = 1. Data are presented as mean ± SEM. **L)** Comparison of maximum VASP fold increase (unpaired t test).

To probe the contributions of each factor to this emergent behavior, we subtracted them individually and monitored F-actin dynamics (Figures 5C and S6C). In the absence of profilin-G-actin, no bundle thickening was observed, demonstrating this occurs due to F-actin polymerization. Omitting VASP produced thinner bundles, supporting a role in stimulating F-actin polymerization, while omitting ɑ-actinin produced mesh-like networks rather than thick bundles. This suggests VASP stimulated polymerization and ɑ-actinin mediated crosslinking are both required to produce well-aligned bundles. Omitting zyxin had only a modest effect under these conditions, suggesting that its interactions with actin regulatory machinery are dispensable for bundle formation and stability in the absence of motor-evoked damage.

We next sought to develop a reconstitution mimicking the active force generating dynamics of stress fibers. A recent report that myosin Va produced beating dynamics in a simpler reconstituted bundle system^40^ motivated us to incorporate soluble myosin VI into our preparation in addition to the coverslip-immobilized pool. This resulted in the formation of bundle networks which hyper-contract within seconds of ATP addition (Figure 5D), producing ruptures in a subset of bundles and the concomitant formation of zyxin patches and bridges (Figure S6D). Eventually, these networks reach a mechanical steady-state where the bundles continue thickening but no longer feature obvious contractile dynamics. When disrupted by laser ablation during this steady-state phase, the severed ends of bundles immediately retract, suggesting networks remain under tension. As has previously been reported in ablation studies of stress fibers in cells, this also results in the rapid recruitment of zyxin to the strain site, reminiscent of zyxin flashes observed at SFSS (Figures 5E-F). We furthermore observed the recovery of actin at these flashes (Figure 5E), suggesting we had minimally reconstituted zyxin-mediated repair of contractile bundles.

The efficient production of strain sites by laser ablation allowed us to dissect the molecular determinants of the repair reaction. As previously reported at SFSS in cells, a subset of flashes failed to produce actin recovery, leading to rupture of the associated bundle (Figure S6E). Omitting either VASP or ɑ-actinin resulted in a significant increase in the fraction of strain sites that failed to repair (Figure S6F), suggesting that both factors are essential in our minimal system. We next analyzed zyxin’s contribution by comparing preparations featuring zyxin^WT^ or zyxin^LDO^ (Figure 5G). We found that zyxin^WT^ flashes repaired significantly more frequently than zyxin^LDO^ flashes. In most of zyxin^LDO^ flashes, the strain site fails to repair and eventually ruptures. This suggests that in the presence of contractile dynamics, binding interactions with VASP and ɑ-actinin coordinated through zyxin’s N-terminus are essential to orchestrate repair. Collectively, these data show that a minimal mixture of zyxin, VASP, ɑ-actinin, profilin-G-actin, and myosin are sufficient to generate contractile bundle networks from an F-actin template and repair them, mimicking stress fiber dynamics in cells.

### Actin restoration initiates at core of strain sites *in vitro* and in cells

We next sought to determine the mechanisms driving actin recovery at strain sites. Similar to the dynamics reported at cellular SFSS^17^, zyxin and VASP are rapidly recruited to reconstituted strain sites, followed by slower recovery of actin signal (Figure 5H). Preparations featuring zyxin^LDO^ instead of zyxin^WT^ displayed minimal actin recovery (Figures 5I-J). We nevertheless observed a moderate degree of VASP recruitment to strain sites in the zyxin^LDO^ condition, which we speculate is mediated by VASP binding to free actin-plus ends exposed by laser ablation. Actin restoration nevertheless fails to occur under these conditions, suggesting VASP engagement of pre-existing actin filaments alone is insufficient to support repair (Figure 5K). Consistently, we observed a significantly higher enrichment of VASP on zyxin^WT^ versus zyxin^LDO^ flashes (Figures 5K-L). Furthermore, in the subset of zyxin^WT^ strain sites which resolved by rupture rather than repair, we observed lower VASP enrichment and a concomitant lack of actin signal recovery (Figures S6G-H), supporting a specific role for zyxin–VASP binding in mediating repair.

The minimal fluorescence bleaching and high signal-to-noise afforded by TIRF imaging allowed us to monitor actin dynamics in reconstituted strain sites with high spatiotemporal resolution (2 seconds / frame). In the zyxin^WT^ condition, we unexpectedly found that repaired strain sites first feature the appearance of increasing actin signal in the core of the zyxin flash, where pre-existing actin signal is minimal, which then spreads outwards to restore the bundle (Figure 6A). In experiments performed without VASP, a small subset of strain sites proceeded towards recovery (Figure S6F). Within this subset, actin density instead weakly recovers inwards from the edges of the strain site, consistent with elongation from pre-existing filament ends in the absence of VASP-mediated stimulation of *de novo* actin polymerization (Figure S7A). Together with our finding that VASP-enriched zyxin patches drive actin nucleation (Figure 4), these results support a model in which reconstituted strain sites are primarily restored through zyxin-mediated amplification of actin nucleation by VASP, initiating at the core of strain sites where cytoskeletal integrity is compromised.

**Figure 6.**
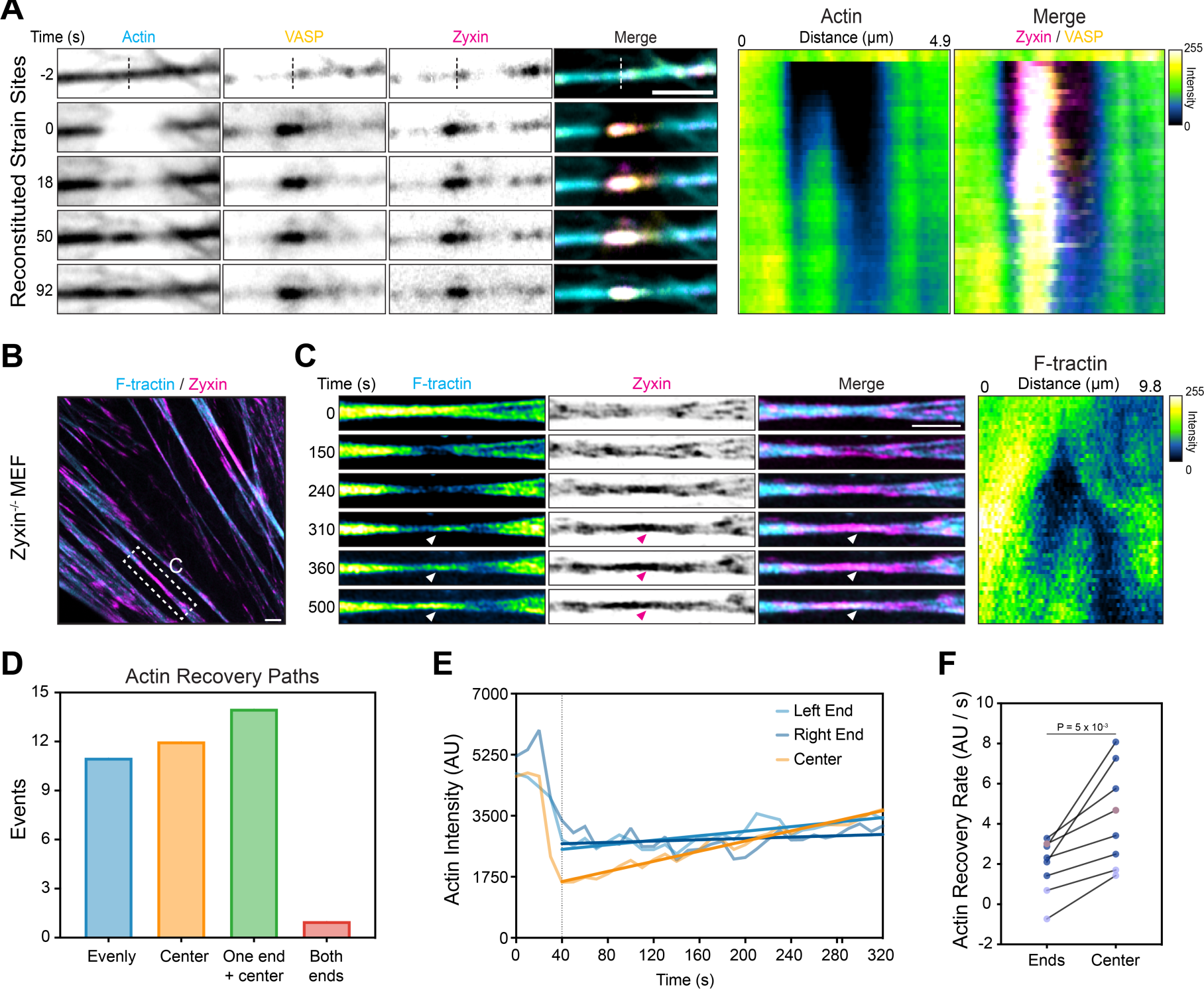
Stress fiber repair initiates within the core of strain sites. **A)** Montage (left) and kymograph (right) of a reconstituted strain site featuring actin recovery that initiates in the core of the zyxin flash. Vertical dashed line indicates site of ablation, which was performed at time = 0 s. Time labels correspond to both montage snapshots and kymograph. Assay was performed with 100 nM zyxin-Halo:JF646, 50 nM VASP-Halo:JF549, 100 nM ɑ-actinin-Halo, 1 μM 30% ATTO-488 labeled G-actin, and 4 μM profilin. Scale bar, 2 μm. **B)** Airyscan image of zyxin^−/−^ mouse embryonic fibroblast expressing F-tractin-mScarlet (cyan) and zyxin-mNeonGreen (magenta). Dashed box highlights SFSS elicited by laser ablation. Scale bar, 2 μm. **C)** Montage (left) and kymograph (right) of SFSS in B. Arrowheads highlight actin recovery initiating in the center of the zyxin flash. Time labels correspond to both montage snapshots and kymograph. Scale bar, 2 μm. **D)** Quantification of actin recovery paths observed at SFSS; n = 19 cells from N = 3 biological replicates. **E)** Quantification of actin recovery over time in indicated regions of a SFSS where recovery was observed to initiate at the center. Vertical dashed line indicates time = 40 s, when SFSS formation initiated. Solid lines represent linear regressions of recovery rate, fit between 40 s and 320 s. **F)** Comparison of actin recovery rates between SFSS center and ends at SFSS where recovery initiated in the center. n = 8 SFSS from N = 3 biological replicates, compared by paired t test.

We next examined whether this mechanism also mediates cytoskeletal repair at SFSS in cells. To achieve sufficient spatiotemporal resolution, we performed live-cell Airyscan imaging of zyxin^−/−^ mouse embryonic fibroblasts (MEFs)^41^ co-expressing zyxin-mNeonGreen and F-tractin-mScarlet while eliciting SFSSs by laser ablation (Figure 6B). Similar to our reconstitution experiments, we observed clear examples of actin density restoration initiating at the core of cellular SFSSs (Figure 6C).To quantify SFSS actin recovery paths, we binned them into four categories: actin recovering evenly across the SFSS (evenly), primarily from the center (center), from one end and the center (one end + center), and inwards from both edges of the SFSS (both ends, Figures 6D and S7B). We observed only one clear example of recovery from both ends, suggesting the vast majority of SFFS repair events feature a component of recovery from the center. Of events where actin recovery initiated primarily from the SFSS core, actin signal increases significantly faster in the core than at the SFSS edges, consistent with a nucleation-mediated mechanism predominating (Figures 6E-F, S7C). The presence of repair path variability suggests that VASP mediated nucleation can be distributed throughout a zyxin flash, rather than being confined to a discrete location, supporting rapid restoration of these micron-scale stress fiber weak points. In summary, our data suggest that force-dependent zyxin assemblies specify VASP’s nucleation of actin filaments at contractile actin-myosin bundle weak points, a mechanism which serves as the dominant actin repair pathway both *in vitro* and in cells.

## DISCUSSION

Here, we demonstrate that a cytoskeletal mechanotransduction pathway, zyxin-dependent repair of mechanically damaged contractile actin-myosin bundles, can be functionally reconstituted with purified proteins. Our data support a model in which force-dependent zyxin assemblies sustain broken actin filaments, templating the formation of multivalent supramolecular assemblies featuring VASP and ɑ-actinin actin regulatory machinery (Figure 7). This local concentration of VASP promotes *de novo* F-actin nucleation and polymerization, which is crosslinked by ɑ-actinin to rebuild stress fibers at the F-actin depleted cores of strain sites. This effectively prioritizes the weakest points in these bundle networks for mechanical restoration.

**Figure 7.**
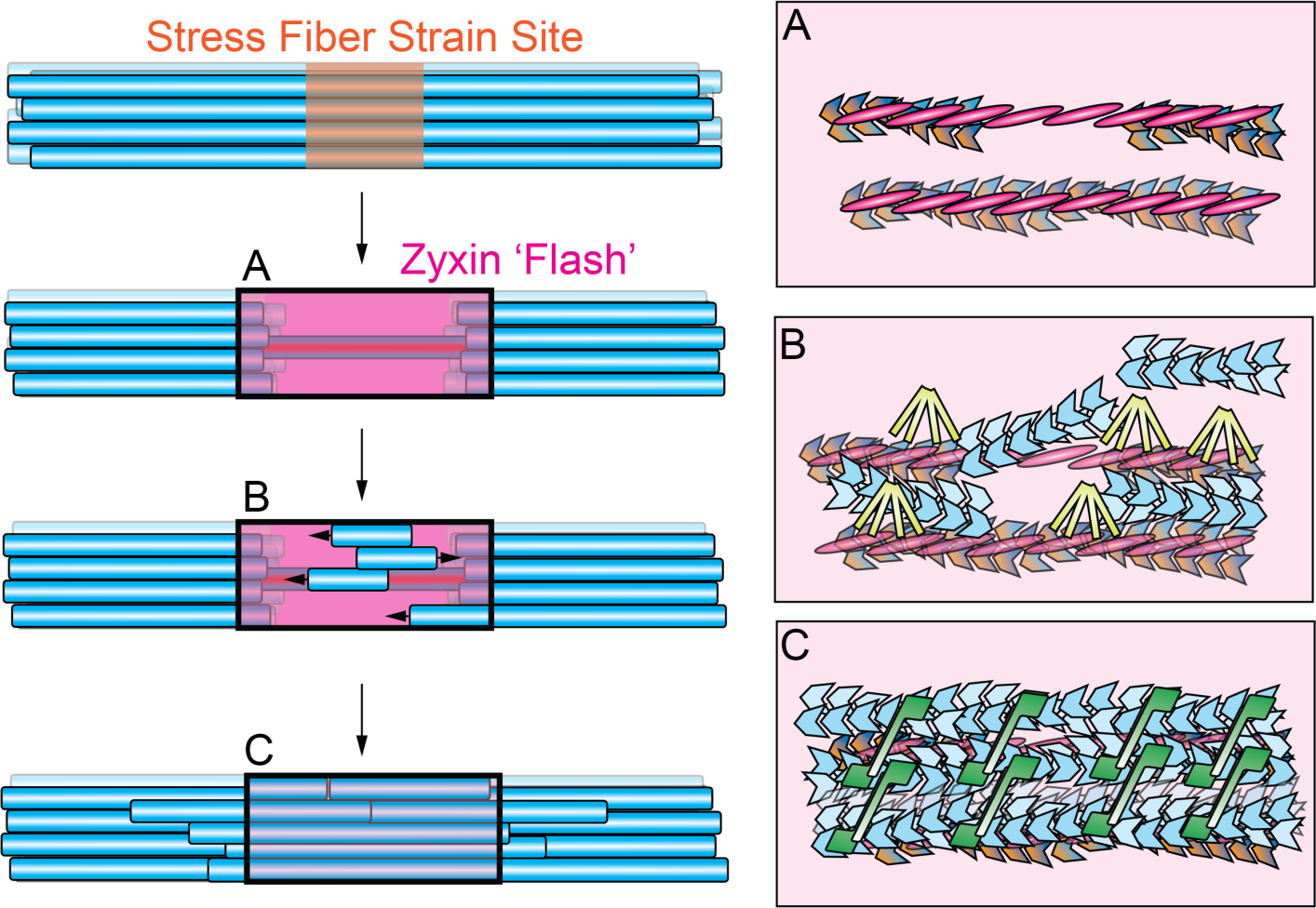
Conceptual model of zyxin-mediated stress fiber repair. Left panel illustrates overall network-scale repair, while right boxes display detailed views of protein components. Actin is displayed in blue; zyxin, magenta; VASP, yellow; ɑ-actinin, green. Strain site formation and concomitant local F-actin depletion results in the formation of force-dependent zyxin assemblies, which may include both actin-bound patches and bridges. These assemblies recruit VASP through multi-valent interactions mediated by zyxin’s N-terminus, which nucleates new F-actin in the strain site core. This F-actin is crosslinked into aligned bundles by ɑ-actinin, which can also engage pre-existing F-actin remnants to restore stress fiber integrity.

While our data are compatible with a “fill-in” mechanism operating in parallel, the network-level mechanism for stress fiber repair we report here offers several potential functional advantages. First, zyxin’s capacity to recruit and concentrate VASP independently of VASP pre-associating with F-actin avoids local competition between VASP productively elongating broken F-actin versus non-productively engaging adjacent intact F-actin at the SFSS periphery. Second, focusing F-actin assembly and crosslinking at SFSS cores facilitates rapid reinforcement of vulnerable filaments bearing extreme loads, thereby preventing catastrophic ruptures. Additionally, while the incorporation of labelled G-actin monomers at SFSS was previously interpreted as F-actin extension from the barbed ends of broken filaments^17^, our data suggest this may have instead represented newly nucleated F-actin. It thus remains possible that broken F-actin at SFSS is rapidly capped in cells, in which case localized assembly of a new F-actin pool would represent an efficient repair mechanism. Future studies examining the detailed spatial composition and nanoscale architecture of SFSS during the repair process will be necessary to fully decipher the interplay of these pathways.

We find that zyxin patches coordinate VASP and ɑ-actinin through two distinct mechanisms to mediate stress fiber repair. VASP’s direct binding to zyxin substantially concentrates the protein, promoting local F-actin assembly in the vicinity of patches. VASP’s role as an F-actin nucleator has been unclear^42–50^, with several studies suggesting that at physiological ionic strength the protein primarily serves as an elongation factor *in vitro*^43–45,51–53^. Conversely, recent work has shown that incorporating VASP into multivalent droplets has the capacity to drive F-actin assembly^49^. VASP-stimulated F-actin nucleation at zyxin patches likely represents a physiological implementation of this phenomenon, although our data do not rule out additional regulatory roles for specific zyxin–VASP binding contacts. On the other hand, zyxin patches do not directly concentrate soluble ɑ-actinin, instead primarily engaging F-actin-bound protein to mediate the reinforcement of mechanically damaged filaments while maintaining stress fiber geometry. Members of the zyxin, paxillin, and FHL family also form patches through their LIM domains^25^, yet they feature distinct accessory domains and suites of binding partners for coordinating downstream mechanotransduction^54^, which likely operate through diverse mechanisms. Beyond its utility for revealing several unanticipated features of the zyxin repair pathway, we anticipate that our reconstitution system’s accessibility to simultaneous TIRF imaging and laser ablation studies will enable future studies dissecting the functional specialization of LIM proteins, as well as the regulation and emergent dynamics of contractile networks more broadly^55^.

To our knowledge, our discovery of LIM protein bridges / tails represents a new class of force-evoked biomolecular assembly. As these assemblies maintain a highly elongated geometry in the absence of underlying F-actin, we speculate they represent force-stabilized polymers of LIM domains. Regardless of their precise molecular architecture, our data strongly imply the existence of both force-dependent LIM–F-actin and LIM–LIM binding interactions, and future studies focusing on the structural basis and detailed biophysical properties of these interfaces will be important for dissecting their interplay. Our finding that bridge formation is a conserved property of force-activated actin-binding LIM domains, as well as previous observations that multiple LIM proteins colocalize at SFSS in cells^16,25,26^, also suggests the intriguing possibility that heterotypic LIM assemblies may form. We speculate these could serve as multi-functional mechanotransduction hubs along stress fibers, in addition to mediating their direct mechanical stabilization, consistent with prior studies highlighting the modularity of LIM domains in mechanical signaling networks^25,26,56^.

Our study also highlights how zyxin coordinates force-dependent binding with multiple classes of multivalent interactions to orchestrate cytoskeletal mechanical homeostasis. While our data support a clear role for multivalency in driving VASP’s rapid localization to SFSS through zyxin’s N-terminus, the precise mechanisms mediating ɑ-actinin’s delayed enrichment remain less clear, as we observed only modest recruitment of soluble ɑ-actinin to zyxin patches along single filaments. Formin mDia1-mediated SF elongation^57^ could push F-actin-bound ɑ-actinin into SFSS via polymerization forces, providing an additional pool of undamaged F-actin available for crosslinking. Alternatively, recruitment could be reinforced through VASP-mediated multivalent binding interactions, as we observed in our droplet experiments. Beyond ɑ-actinin engagement, whether these mechanically-elicited supramolecular assemblies feature emergent biochemical or biophysical properties^58^ that fulfill additional functional roles, e.g. in strain site stabilization or specifying the local architecture of F-actin networks as they are rebuilt, remain important topics for future studies.

## Acknowledgements

We thank M. Beckerle for the gift of zyxin^−/−^ MEFs, as well as M. Smith, M. Beckerle, K. Baboolall, K. Landy, D. Kovar, and M. Gardel for discussion of unpublished data. We gratefully acknowledge use of the Rockefeller University Bio-Imaging Resource Center (BIRC) for iSIM, ringTIRF, and Airyscan confocal imaging, and P. Banerjee, V. Sharma, C. Pyrgaki, and A. North for microscopy assistance. This work was funded by a National Institute of General Medical Sciences of the National Institutes of Health (NIH) grant (R01GM146880) to G.M.A. D.Y.Z.P was supported by a National Heart, Lung, and Blood Institute of the NIH fellowship (F31HL165906).

## Resource Availability

All resources and reagents reported in this study are freely available, and requests should be directed to the lead contact, Gregory M. Alushin.

## Data Availability

All imaging data will be made available at Zenodo (XXXX). All other data are presented in the manuscript.

## Code Availability

Custom code will be made available at https://www.github.com/alushinlab/zyxin-repair as open source.

## Contributions

D.Y.Z.P. and G.M.A. conceived the project and designed research. D.Y.Z.P. performed all experiments with zyxin and all data analyses. X.S. performed experiments with FHL3 and assisted with VASP binding analyses. D.Y.Z.P. and G.M.A. wrote the paper with input from X.S.

## VIDEO LEGENDS

**Video S1. LIM protein bridges tether broken F-actin fragments. Related to Figure 1**.

Representative movies of LIM protein F-actin-bound patches, bridges, and tails. Zyxin’s LIM domains are sufficient to form bridges. FHL3, another mechanosensitive LIM protein, can also form bridges. Scale bar, 2 μm. Time is minutes : seconds.

**Video S2. VASP dynamics on zyxin patches. Related to Figure 2**.

Representative movies of VASP binding to zyxin patches. Scale bar, 2 μm. Time is minutes : seconds.

**Video S3. ɑ-actinin–zyxin binding and F-actin bundling dynamics. Related to Figure 3**.

Representative movies of ɑ-actinin binding to zyxin patches and clusters in the presence and absence of force (+/− ATP). Zyxin patches serve as a molecular guide for ɑ-actinin-mediated zippering of translocating actin filaments into aligned bundles. Scale bar, 2 μm. Time is minutes : seconds.

**Video S4. Dynamics of Zyxin, VASP, and ɑ-actinin droplets. Related to Figure S4.**

Representative movies of droplet fusion and fluorescence recovery after photobleaching (FRAP). Scale bar, 2 μm. Time is minutes : seconds.

**Video S5. VASP-mediated F-actin nucleation and polymerization-coupled extrusion at zyxin patches in the presence of myosin forces. Related to Figure 4**.

Representative movies of immobilized myosin V and myosin VI conditions. Scale bar, 2 μm. Time is minutes : seconds.

**Video S6. Dynamics of reconstituted actin-myosin bundles and zyxin-mediated repair. Related to Figure 5**.

Representative movies of reconstituted non-contractile and contractile bundles, including examples of spontaneous zyxin bridge formation, zyxin flashes and zyxin mediated repair, as well as catastrophic bundle rupture at a zyxin flash. Scale bar, 2 μm. Time is minutes : seconds.

**Video S7. Dynamics of F-actin recovery paths at strain sites. Related to Figure 6**.

Representative movies of F-actin recovery initiating in the cores of both reconstituted strain sites and SFSS of zyxin^−/−^ MEFs expressing zyxin-mNeonGreen and F-tractin-mScarlet, followed by examples of all F-actin recovery paths observed in cells. Scale bar, 2 μm. Time is minutes : seconds.

---

**Figure S1.**
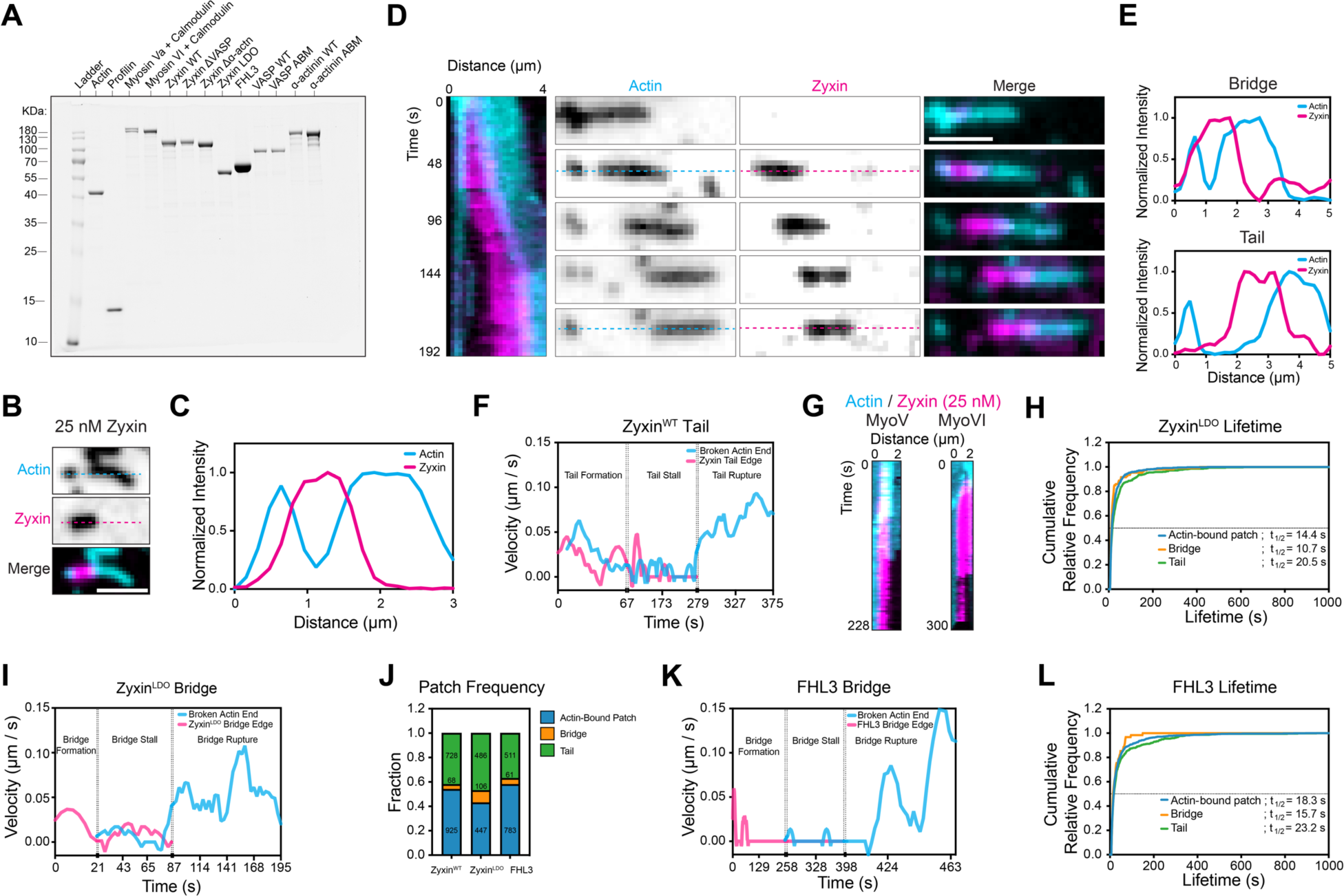
Zyxin bridges and tails feature similar architectures and dynamics. Related to Figure 1. **A)** Coomassie-stained SDS-PAGE of purified proteins. **B)** TIRF snapshot of a zyxin bridge formed in the presence of 100% ATTO 488-labeled actin filaments (cyan) and 25 nM zyxin-Halo:JF646 (magenta). Scale bar, 2 μm. **C)** Normalized fluorescence intensity along dashed lines shown in B. **D)** Kymograph (left) and montage (right) of a zyxin bridge forming in the presence of 250 nM zyxin-Halo:JF646, then converting into a tail. Time labels correspond to both montage snapshots and kymograph. Scale bar, 2 μm. **E)** Normalized fluorescence intensity along dashed lines shown in D (bridge, 48 s; tail, 192 s). **F)** Instantaneous velocity over time of the interface between a zyxin tail and a broken F-actin fragment. **G)** Kymographs of zyxin bridges converting into tails in the presence of myosin Va alone (left) or myosin VI alone (right). Zyxin-Halo:JF646, 25 nm. **H)** Cumulative relative frequency of zyxin LDO actin-bound patch, bridge, and tail lifetimes. Dotted line = 0.5. **I)** Instantaneous velocity over time of the interface between a zyxin LDO bridge and a broken F-actin fragment. **J)** Patch category frequency across LIM proteins. Number of each structure observed in each category are indicated; N ≥ 2 biological replicates. **K)** Instantaneous velocity over time of the interface between a FHL3 bridge and a broken F-actin fragment. **L)** Cumulative relative frequency of FHL3 actin-bound patch, bridge, and tail lifetimes. Dotted line = 0.5.

**Figure S2.**
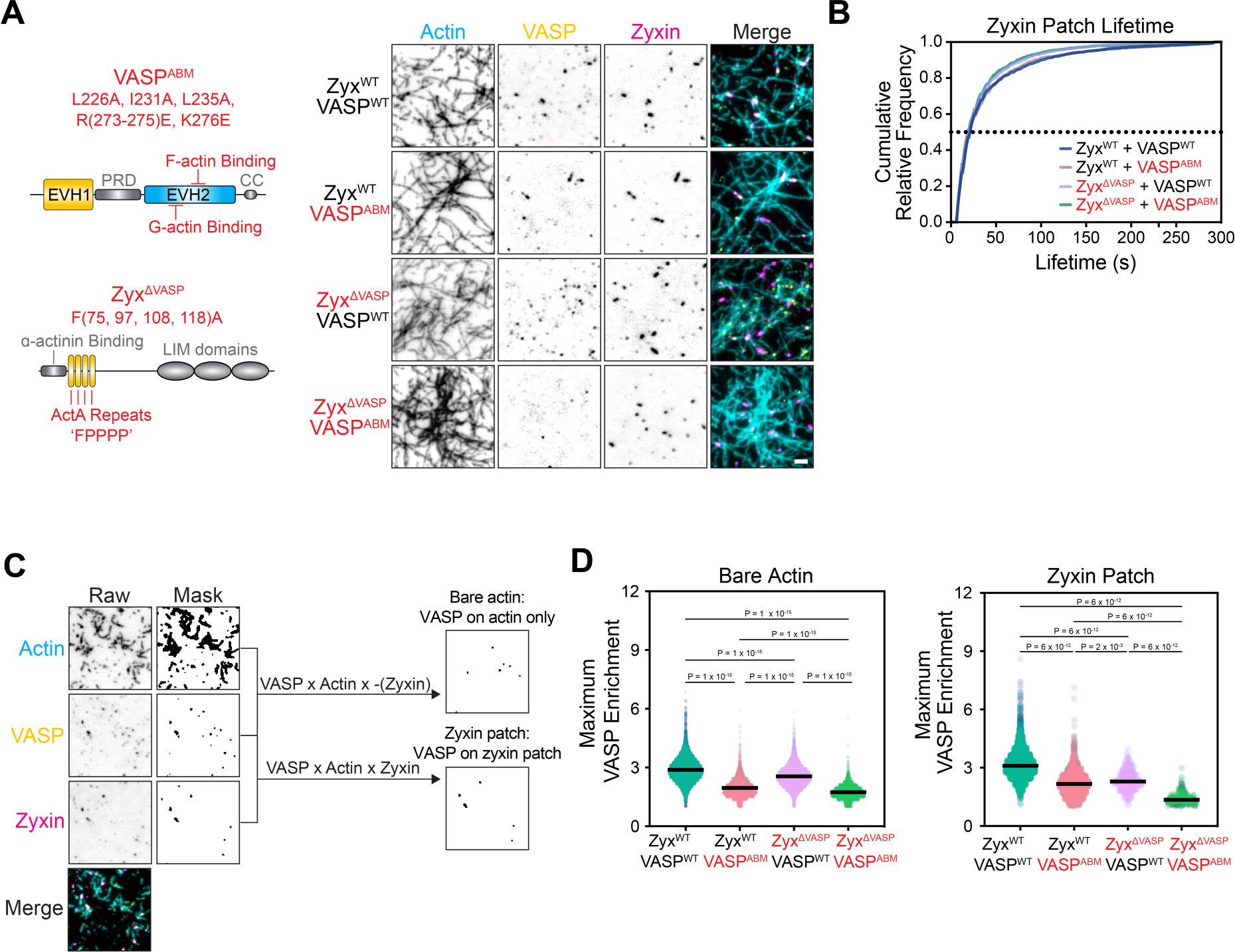
Additional analysis of VASP enrichment on zyxin patches. Related to Figure 2. **A)** Left: Schematic of zyxin^ΔVASP^ and VASP^ABM^ mutant constructs. Right: TIRF snapshots of force reconstitution assays featuring the indicated constructs, performed with 30% ATTO 488-labeled actin filaments (cyan), 50 nM VASP-Halo:JF549 (yellow), and 25 nM zyxin-Halo:JF646 (magenta). Scale bar, 2 μm. **B)** Cumulative relative frequency of zyxin patch lifetimes in the presence of the indicated constructs. Dotted line = 0.5. **C)** Workflow for quantifying VASP on bare F-actin vs. actin-bound zyxin patches. **D)** Maximum VASP enrichment on bare F-actin (left) or zyxin patches (right) in the presence of the indicated constructs. Bars represent means; 367 ≤ n ≤ 4479 from N = 2 biological replicates (represented by shades). Conditions were compared by Turkey’s multiple comparisons test after ordinary one-way ANOVA.

**Figure S3.**
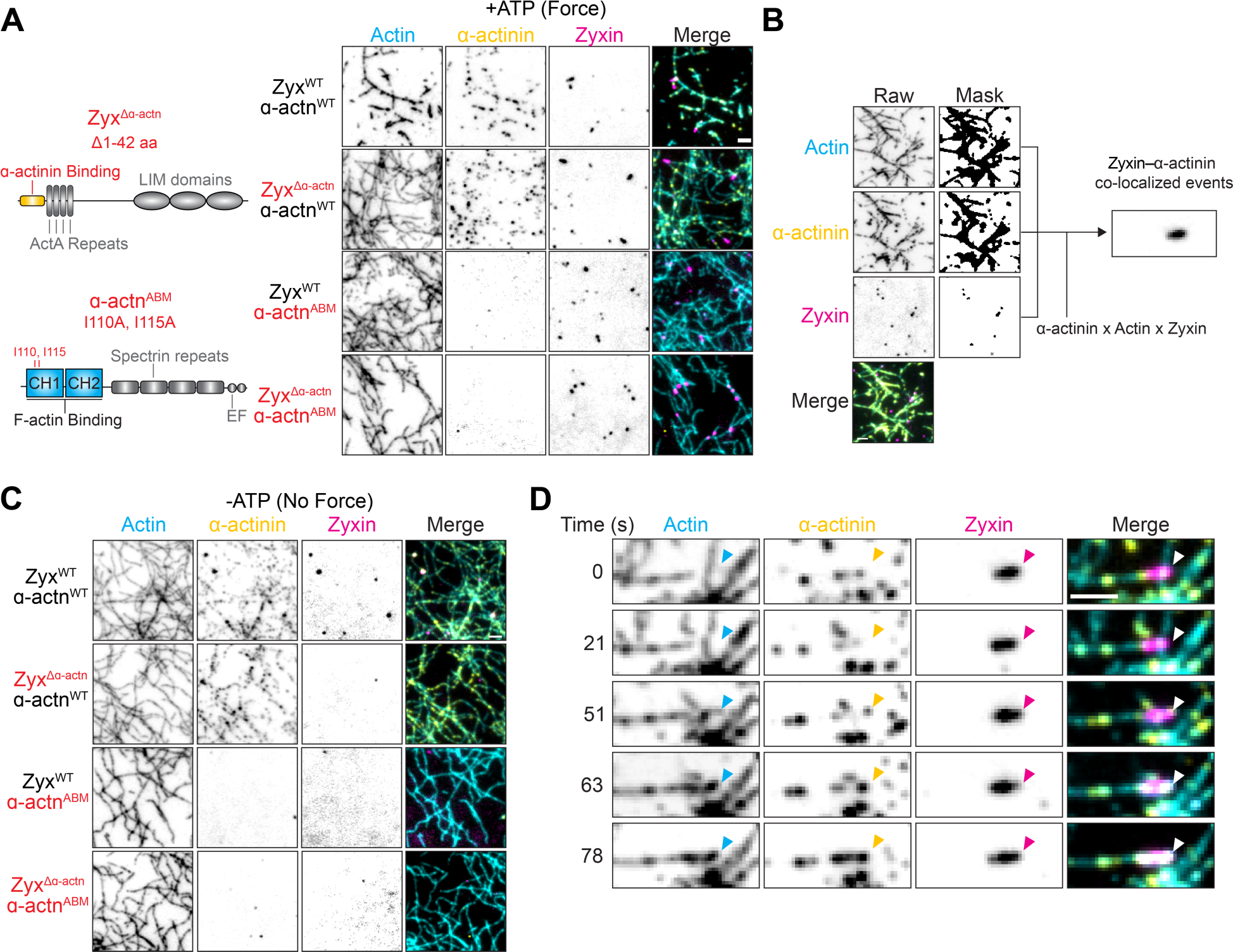
F-actin bound ɑ-actinin weakly engages zyxin patches. Related to Figure 3. **A)** Left: Schematic of zyxin^Δɑ-actinin^ and ɑ-actinin^ABM^ mutant constructs. Right: TIRF snapshots of force reconstitution assays performed with 30% ATTO 488-labeled actin filaments (cyan), 50 nM ɑ-actinin-Halo:JF549 (yellow), and 25 nM zyxin-Halo:JF646 (magenta) in the presence of the indicated constructs and immobilized myosin forces (+ATP). Scale bar, 2 μm. **B)** Workflow for quantifying zyxin–ɑ-actinin colocalization events. **C)** TIRF snapshots of reconstitution assays performed as in A, but in the absence of force generation (-ATP). Scale bar, 2 μm. **D)** Montage of multiple actin filaments (cyan) bundling along a zyxin patch (magenta) through ɑ-actinin clusters (yellow). Patch position is indicated with arrowheads. The assay was performed with 30% ATTO 488-labeled actin filaments, 50 nM ɑ-actinin-Halo:JF549, and 25 nM zyxin-Halo:JF646. Scale bar, 2 μm.

**Figure S4.**
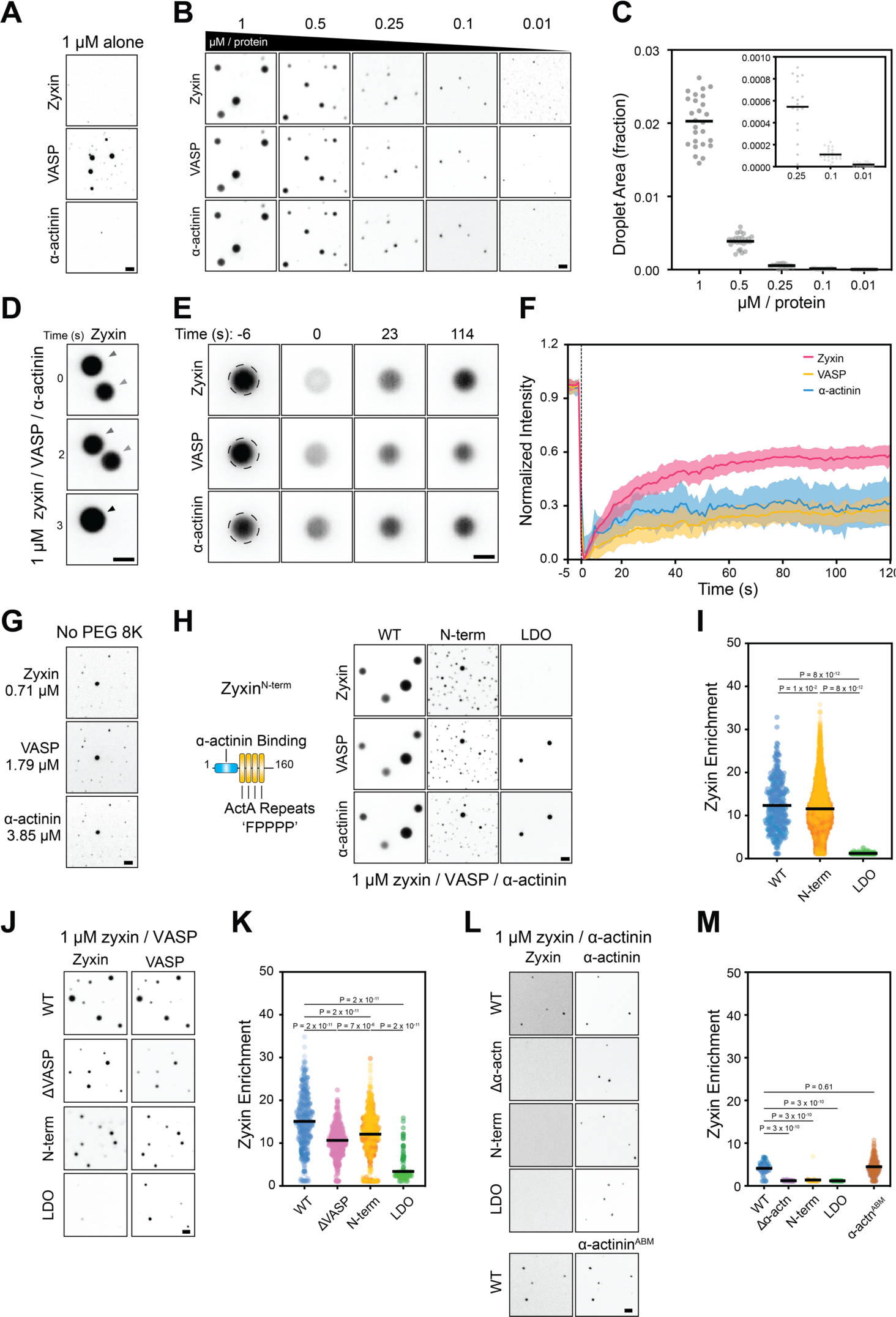
Zyxin, VASP, and ɑ-actinin form droplets through multivalent interactions. Related to Figure 5. **A)** iSIM snapshots of 1 μM zyxin-Halo:JF646 (top), 1 μM VASP-Halo:JF549 (middle), and 1 μM ɑ-actinin-Halo:AF488 (bottom), individually in the presence of 3% PEG 8K. Scale bar, 2 μm. **B)** iSIM snapshots of tripartite droplets across varying concentrations of zyxin-Halo:JF646, VASP-Halo:JF549, and ɑ-actinin-Halo:AF488 in the presence of 3% PEG 8K. Scale bar, 2 μm. **C)** Quantification of droplet size distribution (as fraction of field of view) from experiments presented in B. n = 12 fields from N = 2 biological replicates. **D)** Montage of two droplets undergoing fusion. Tripartite droplets were assembled from 1 μM zyxin-Halo:JF646, 1 μM VASP-Halo:JF549, and 1 μM ɑ-actinin-Halo:AF488 in the presence of 3% PEG 8K. Scale bar, 2 μm. **E)** Montage of fluorescence recovery after photobleaching (FRAP) of a tripartite droplet prepared as in D. Dashed circle represents the region illuminated at time = 0 s. Scale bar, 2 μm. **F)** Quantification of experiments presented in E. Dashed line represents time = 0 s. Data are presented as mean ± SD; n = 14 droplets bleached from N = 2 biological replicates. **G)** iSIM snapshot of tripartite droplets formed with indicated physiological concentrations of zyxin-Halo:JF646, VASP-Halo:JF549, and ɑ-actinin-Halo:AF488 in the absence of PEG 8K. Scale bar, 2 μm. **H)** Left: Schematic of zyxin^Nterm^ construct. Right: iSIM snapshots of tripartite droplets formed with indicated zyxin constructs in the presence of 3% PEG 8K. Samples were prepared with 1 μM zyxin-Halo:JF646, 1 μM VASP-Halo:JF549, and 1 μM ɑ-actinin-Halo:AF488. Scale bar, 2 μm. **I)** Quantification of zyxin enrichment in droplets from experiments presented in H. Bars represent means; 338 ≤ n ≤ 5103 from N = 2 biological replicates (represented by shades). **J)** iSIM snapshots of bipartite zyxin and VASP droplets formed with indicated zyxin constructs in the presence of 3% PEG 8K. Samples were prepared with 1 μM zyxin-Halo:JF646 and 1 μM VASP-Halo:JF549. Scale bar, 2 μm. **K)** Quantification of zyxin enrichment in droplets from experiments presented in J. Bars represent means; 203 ≤ n ≤ 651 from N = 2 biological replicates (represented by shades). **L)** iSIM snapshots of bipartite zyxin and ɑ-actinin droplets formed with indicated zyxin and ɑ-actinin constructs in the presence of 3% PEG 8K. Samples were prepared with 1 μM zyxin-Halo:JF646 and 1 μM ɑ-actinin-Halo:AF488. Scale bar, 2 μm. **M)** Quantification of zyxin enrichment in droplets from experiments presented in L. Bars represent means; 34 ≤ n ≤ 359 from N = 2 biological replicates (represented by shades). All statistical comparisons were performed with Turkey’s multiple comparisons test after one-way ordinary ANOVA.

**Figure S5.**
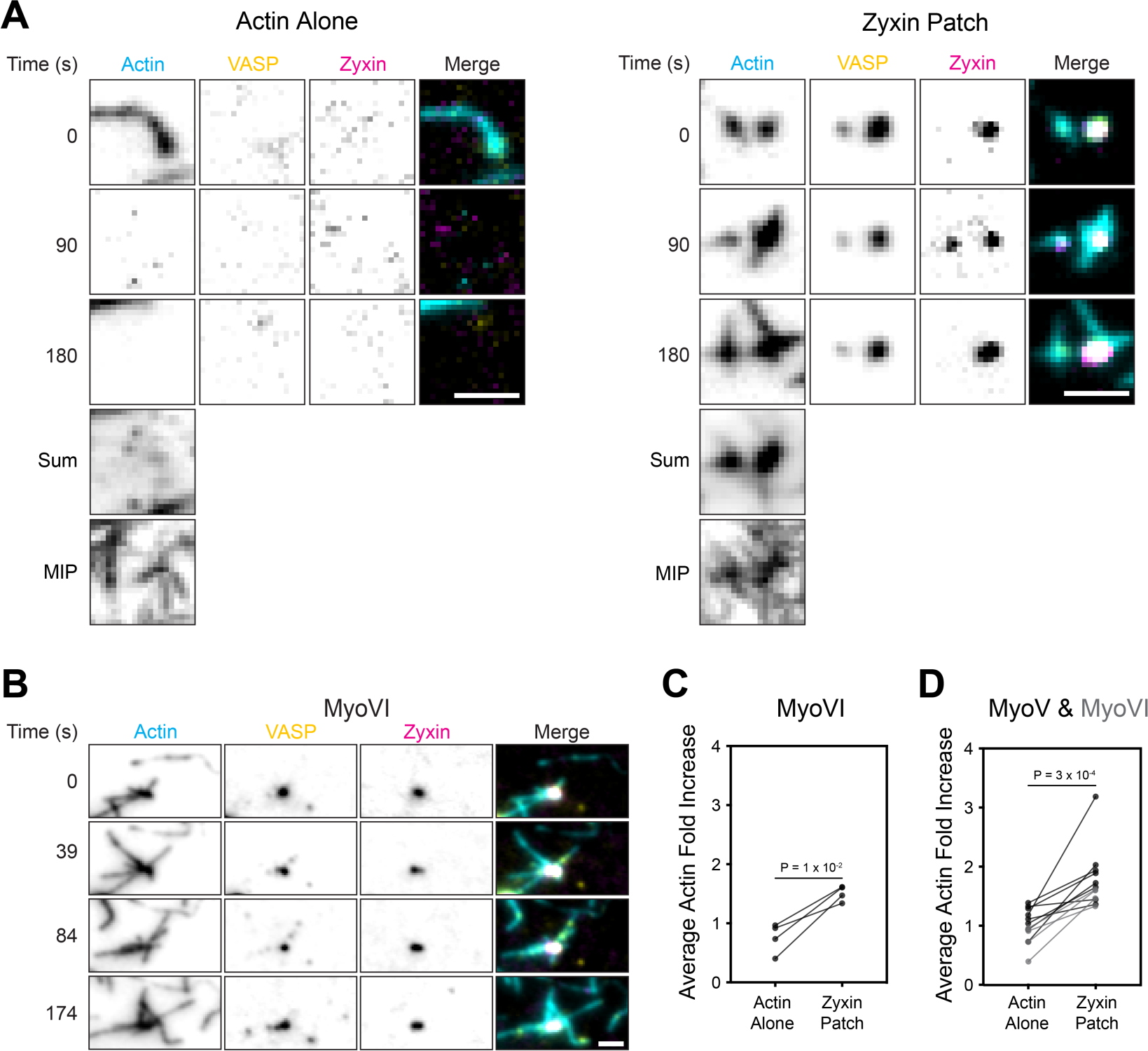
Zyxin patches promote VASP-mediated actin nucleation and polymerization. Related to Figure 4. **A)** Montages of undetectable actin filament (cyan, 30% ATTO 488-labeled) nucleation and polymerization at a bare F-actin region (left) versus at a VASP-enriched (yellow) zyxin patch (magenta, right) in the presence of immobilized myosin VI forces (+ATP). Assay was performed with 0.5 μM 30% ATTO-488 G-actin, 2 μM profilin, 50 nM VASP-Halo:JF549, and 25 nM zyxin-Halo:JF646. Scale bar, 2 μm. **B)** Montage of F-actin nucleation and polymerization-coupled extrusion from a zyxin patch formed in the presence of myosin VI (+ATP). Assay conditions were otherwise identical to Figure 4B. **C)** Pairwise comparison of the average actin intensity fold increase at VASP-enriched zyxin patches versus bare F-actin regions in the presence of myosin VI forces (+ATP). Assays were performed as in A. n = 4 trials (imaging 1-5 field of view per trial) from N = 2 biological replicates, compared by paired t test. **D)** Pooled paired comparisons of average actin fold increase at VASP-enriched zyxin patches vs. bare F-actin across trials, in the presence of either immobilized myosin V (black) or myosin VI (grey) individually. n = 12 trials from N = 2 biological replicates, compared by paired t test.

**Figure S6.**
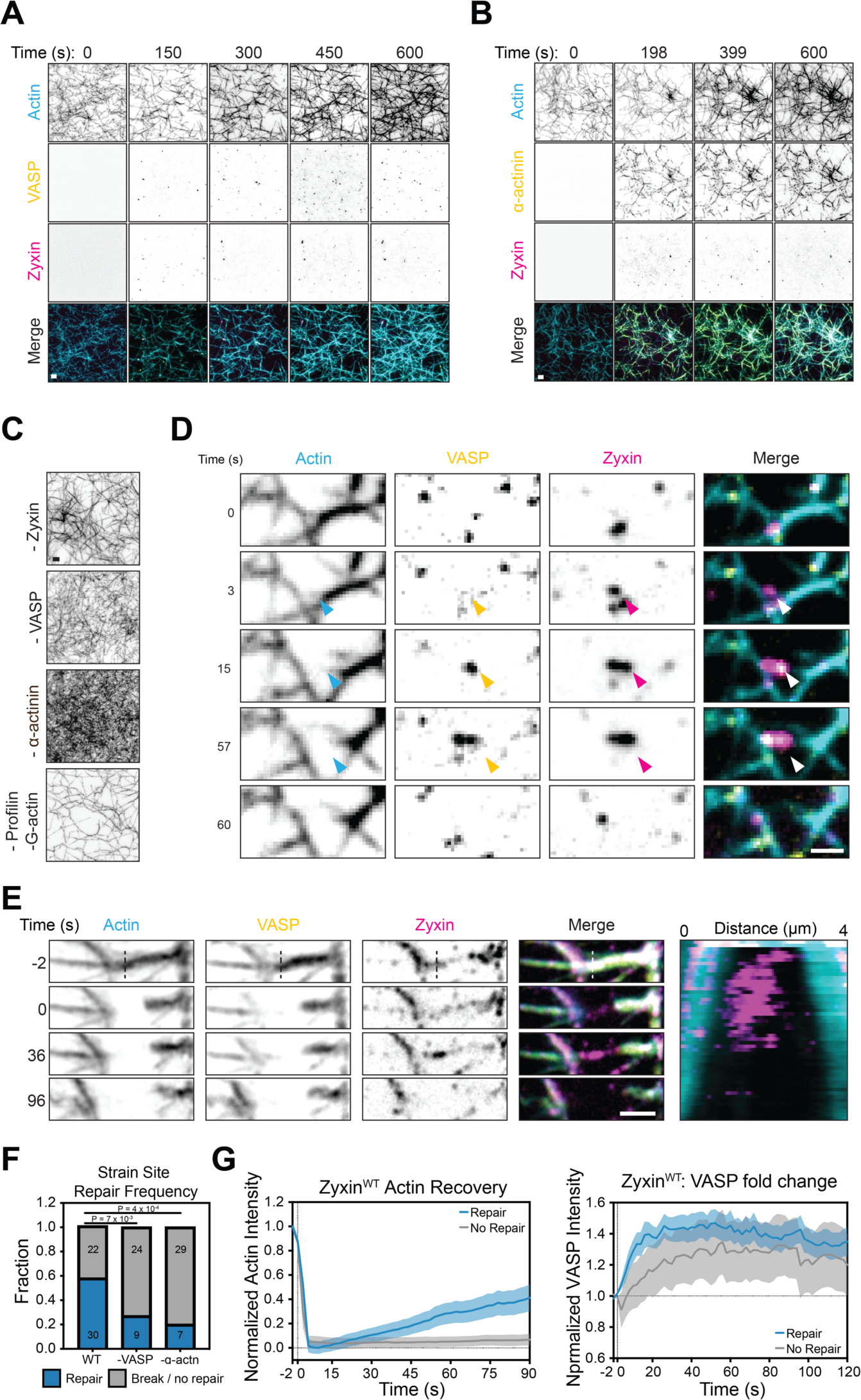
Further characterization of reconstituted contractile bundles and F-actin repair. Related to Figure 5. **A)** Montage of bundle reconstitution assay featuring labeled VASP and unlabeled ɑ-actinin. Assay was prepared with 30% ATTO 488-labeled F-actin (cyan), 25 nM zyxin-Halo:JF646 (magenta), 50 nM VASP-Halo:JF549 (yellow), 50 nM ɑ-actinin-Halo (unlabeled), 0.5 μM 30% ATTO-488 labeled G-actin, and 2 μM profilin. Scale bar, 2 μm. **B)** Montage of bundle reconstitution assay featuring labeled ɑ-actinin and unlabeled VASP. Assay was prepared with 30% ATTO 488-labeled F-actin (cyan), 25 nM zyxin-Halo:JF646 (magenta), 50 nM VASP-Halo:JF549 (unlabeled), 50 nM ɑ-actinin-Halo (yellow), 0.5 μM 30% ATTO-488 labeled G-actin, and 2 μM profilin. Scale bar, 2 μm. **C)** Representative TIRF snapshots of bundle reconstitutions at time = 600 s where indicated components were omitted. Scale bar, 2 μm. **D)** Montage of a zyxin bridge (magenta arrowhead) which forms within a contractile network and accumulates VASP (yellow arrowhead), then ruptures. Assay was prepared with 100 nM zyxin-Halo:JF646, 50 nM VASP-Halo:JF549, 100 nM ɑ-actinin-Halo, 1 μM 30% ATTO-488 labeled G-actin, and 4 μM profilin. Scale bar, 2 μm. **E)** Montage (left) and kymograph (right) of a laser ablation-generated strain site that fails to repair and ruptures. Vertical dashed lines indicate site of ablation, performed at time = 0 s. Time labels correspond to both montage snapshots and kymograph. Scale bar, 2 μm. **F)** Quantification of strain site repair frequency in the absence of VASP or ɑ-actinin in assays otherwise performed as in D. Number of strain sites in each category are indicated; N ≥ 3 biological replicates. Conditions were compared by Fisher’s exact test. **G)** Normalized actin fluorescence intensity (left) and VASP intensity fold increase (right) over time at strain sites that either repaired (blue, n = 17) or did not repair (grey, n = 7) in the presence of zyxin^WT^. Vertical dashed lines indicate time = 0 s, when laser ablation was performed. Horizontal dashed line indicates baseline VASP intensity = 1. N = 5 biological replicates. Data are presented as mean ± SEM.

**Figure S7.**
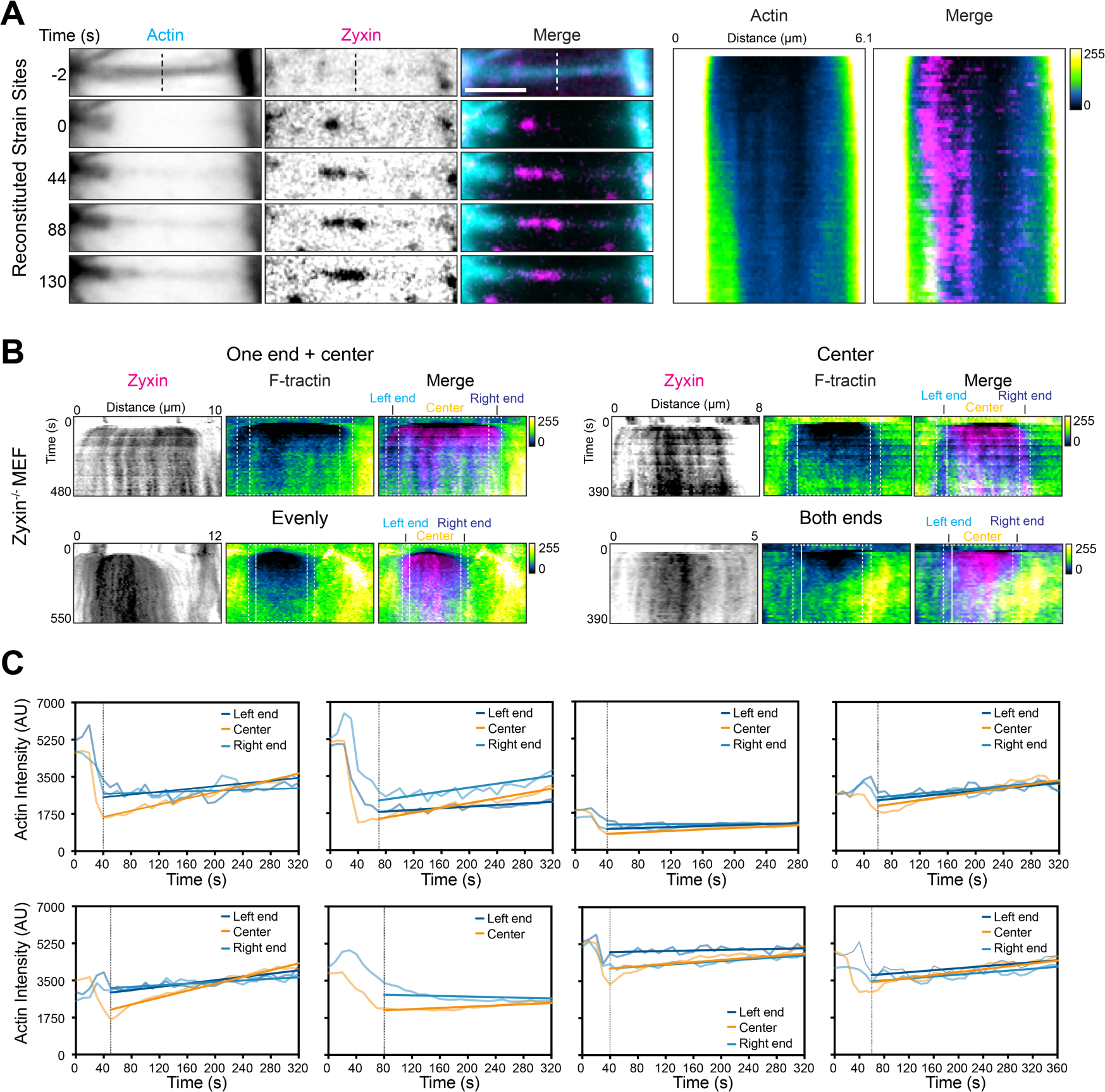
Additional analysis of SFSS actin repair paths. Related to Figure 6. **A)** Montage (left) and kymograph (right) of a reconstituted strain site from a preparation lacking VASP, where actin recovery occurs at the strain site edges. Vertical dashed line indicates site of ablation, which was performed at time = 0 s. Time labels correspond to both montage snapshots and kymograph. Assay was performed with 100 nM zyxin-Halo:JF646, 100 nM ɑ-actinin-Halo, 1 μM 30% ATTO-488 labeled G-actin, and 4 μM profilin. Scale bar, 2 μm. **B)** Kymographs of cellular SFSS representing each actin recovery path quantified in Figure 6D. Dashed boxes highlight quantified regions; each end was assigned as 10% of the overall strain site length. **C)** Actin recovery rate quantification for all cellular SFSS compared in Figure 6F, analyzed as in Figure 6E.

**Table S1:** Component concentrations in force reconstitution assays.

| Force Reconstitution Assays | Zyxin (nM) | VASP (nM) | α-actinin (nM) | Profilin (μM) | G-actin (μM) | ATP (mM) |
| --- | --- | --- | --- | --- | --- | --- |
| Zyxin alone (Fig. 1) | 25 / 250 | 0 | 0 | 0 | 0 | 0.5 |
| Zyxin + VASP (Fig. 2) | 25 | 50 | 0 | 0 | 0 | 0.5 |
| Zyxin + α-actinin (Fig. 3) | 25 / 250 | 0 | 50 | 0 | 0 | 1 |
| Zyxin + VASP + Profilin-G-actin (Fig. 4) | 25 | 50 | 0 | 2 | 0.5 | 2 |
| Zyxin + VASP + α-actinin + Profilin-G-actin (Fig. 5,6) | 25 | 50 | 50 | 2 | 0.5 | 2 |

## METHODS

### Plasmids and cloning

Plasmids were propagated in NEB 5-alpha competent E. coli cells (New Englad Biolabs) and purified with QIAprep spin miniprep kits (Qiagen) or the PureYield plasmid maxiprep system (Promega) before transfection.

Constructs for human myosin Va (amino acids 1-1091), myosin VI (amino acids 1-1021), calmodulin, and FHL3-Halo were reported previously^1^. Human Zyxin-Halo constructs were subcloned into a pCAG mammalian expression vector with a C-terminal 3C protease cleavage site and eGFP tag for affinity purification. Human VASP-Halo constructs were subcloned into a pET15b bacterial expression vector containing an N-terminal Strep-Tag II and 3C cleavage site, as well as a C-terminal 3C site and 6xHis-Tag. Human ɑ-actinin-Halo constructs were subcloned into a pET15b bacterial expression vector containing an N-terminal 6xHis-Tag and Tobacco Etch Virus (TEV) cleavage site. Human profilin in a pMW plasmid was a generous gift from Henry Higgs, Dartmouth University. Human Zyxin-mNeonGreen and F-tractin-mScarlet were subcloned into a pM98 mammalian expression vector. The template for mScarlet was a gift from Dorus Gadella (University of Amsterdam) via Addgene (plasmid #85054). All plasmids were assembled with Gibson cloning, and point mutations were generated by site-directed mutagenesis.

### Protein expression and purification

Human Zyxin, myosin Va, and myosin VI were expressed in HEK293F cells as previously reported^1^. In brief, cultures were transfected at a density of 1.8×10^6^ cells/mL. For 400 mL of cells, 400 μg of plasmid was pre-mixed with 1.2 mL of 1mg/mL PEI MAX (Polysciences) in 15 mL of FreeStyle 293 expression medium. Myosins were co-transfected with calmodulin at a 1:6 DNA mass ratio. After a 20-minute incubation at room temperature, DNA:PEI complexes were added to the cells for transfection. Cells were grown on an orbital shaker in a 37°C incubator at 8% CO_2_ and harvested by centrifugation at 1000 x *g* for 5 minutes 60-72 hours post transfection.

Human VASP, ɑ-actinin, calmodulin, 6XHis-TEV protease, and profilin were expressed in Rosetta2(DE3) E. coli cells (Novagen) grown in LB media at 37°C. 6XHis-3C protease was expressed in BL21(DE3) E. coli cells (Thermo Scientific). After reaching an optical density of 0.6, expression was induced with 0.5 mM Isopropyl ß-D-1-thiogalactopyranoside (IPTG). For expression of VASP and profilin, cells were grown at 16°C for 16-20 hours after induction. For expression of ɑ-actinin, cells continued to grow at 37°C for 6 hours after induction. Cell pellets were collected by centrifugation at 4000 x *g* for 10 minutes, flash frozen in liquid nitrogen, and stored at −80°C until use

FHL3 and zyxin was purified as previously described^1^. Cells were resuspended in Lysis Buffer A (50 mM Tris-HCl pH 8.0, 300 mM NaCl, 0.2% nonyl phenoxypolyethoxylethanol (NP40), 3 mM 2-mercaptoethanol (βME), 2 mM ATP, 1 mM phenylmethylsulfonyl fluoride (PMSF), 1 μg/mL aprotinin, leupeptin, and pepstatin A) and nutated for 30 minutes. The lysate was then clarified by centrifugation for 30 minutes at 48,000 x *g*. The supernatant was mixed with NHS-activated Sepharose 4 Fast Flow resin (GE Healthcare) coupled to an anti-GFP nanobody expressed and purified by published protocols^2^ and incubated on a rocker for 2 hours. The resin was then washed with Lysis Buffer A without detergent and protease inhibitors, then incubated in the presence of 6XHis-3C protease that was purified by published protocols^3^ overnight. The supernatant was collected and purified by anion exchange chromatography with a Mono Q column (Cytiva) followed by size exclusion chromatography (SEC) with a Superdex 200 increase column (Cytiva) in Gel Filtration Buffer (10 mM Tris-HCl pH 8.0, 100 mM NaCl, 3 mM dithiothreitol (DTT)). Peak fractions were pooled, concentrated, flash frozen in liquid nitrogen and stored at −80°C until use. FHL3, Zyxin LDO, and N-term was concentrated with a 30 kDa molecular weight cut off (MWCO) concentrator (Amicon), and all full-length or Δɑ-actinin constructs were concentrated with a 50 kDa MWCO concentrator (Amicon).

To purify VASP, cells were resuspended in Lysis Buffer B (50 mM Tris-HCl pH 8.0, 300 mM NaCl, 20 mM imidazole, 3 mM βME, 2 mM ATP, 1 mM PMSF, 1 μg/mL aprotinin, leupeptin, and pepstatin A) and lysed with an Avestin Emulsiflex C5 homogenizer. The lysate was clarified by centrifugation for 30 minutes at 48,000 x *g*. The supernatant was then applied to a Ni-NTA agarose gravity column (Qiagen). The column was washed with Lysis Buffer B, then eluted with Lysis Buffer B supplemented with 250 mM imidazole without protease inhibitors. The eluant was further purified by a StrepTrap XT column (GE Healthcare). Peak fractions were pooled, then dialyzed (SnakeSkin, 10K MWCO) against Dialysis Buffer A (25 mM Tris-HCl pH 8.0, 300 mM NaCl, 3 mM βME) in the presence of 6XHis-3C protease overnight. The mixture was again applied to a Ni-NTA agarose gravity column to remove 3C protease, and finally purified by SEC with a Superose 6 increase column (Cytiva) in Gel Filtration Buffer. Peak fractions were pooled, concentrated in a 100 kDa MWCO concentrator (Amicon), flash frozen in liquid nitrogen, and stored at −80°C until use.

ɑ-actinin purification was adapted from published protocols^4^. Cells were resuspended in Lysis Buffer B (50 mM Tris-HCl pH 8.0, 300 mM NaCl, 20 mM imidazole, 3 mM βME, 2 mM ATP, 1 mM PMSF, 1 μg/mL aprotinin, leupeptin, and pepstatin A) and lysed with an Avestin Emulsiflex C5 homogenizer. The lysate was clarified by centrifugation for 30 minutes at 48,000 x *g*. The supernatant was then applied to a Ni-NTA agarose gravity column. The column was washed with Lysis Buffer B, then eluted with Lysis Buffer B supplemented with 250 mM Imidazole without protease inhibitors. The eluant was dialyzed (SnakeSkin, 10K MWCO) against Dialysis Buffer A (25 mM Tris-HCl pH 8.0, 300 mM NaCl, 3 mM βME) in the presence of 6XHis-TEV protease that was purified by published protocols^5^ overnight. The mixture was then again applied to a Ni-NTA agarose gravity column to remove TEV protease. The flowthrough was collected and purified by anion exchange chromatography with a HiTrap Q HP column (Cytiva) followed by SEC with a Superose 6 increase column in Gel Filtration Buffer. Peak fractions were pooled, concentrated in a 100 kDa MWCO concentrator (Amicon), flash frozen in liquid nitrogen, and stored at −80°C until use.

To purify calmodulin, cells were resuspended in Lysis Buffer B (50 mM Tris-HCl pH 8.0, 300 mM NaCl, 20 mM imidazole, 3 mM βME, 2 mM ATP, 1 mM PMSF, 1 μg/mL aprotinin, leupeptin, and pepstatin A) and lysed with an Avestin Emulsiflex C5 homogenizer. The lysate was clarified by centrifugation for 30 minutes at 48,000 x *g*. The supernatant was then applied to a Ni-NTA agarose gravity column. The column was washed with Lysis Buffer B, then eluted with Lysis Buffer B supplemented with 250 mM imidazole without protease inhibitors. The eluant was dialyzed (SnakeSkin, 10K MWCO) against Dialysis Buffer A (25 mM Tris-HCl pH 8.0, 300 mM NaCl, 3 mM βME) in the presence of 6XHis-3C protease overnight. The mixture was then applied to a Ni-NTA agarose gravity column to remove 3C protease. The flowthrough was collected and purified by anion exchange chromatography with a HiTrap Q HP column followed by SEC with a Superose 6 increase column in Gel Filtration Buffer. Peak fractions were pooled, concentrated in a 3 kDa MWCO concentrator (Amicon), flash frozen in liquid nitrogen, and stored at −80°C until use.

Myosin Va and VI were purified as previously reported^1^. In brief, cells were resuspended in Lysis Buffer C (50 mM Tris-HCl pH 8.0, 150 mM NaCl, 0.2% CHAPS, 2 mM MgCl_2_, 2 mM ATP, 1 mM PMSF, 1 μg/mL aprotinin, leupeptin, and pepstatin A) and nutated for 30 minutes. The lysate was clarified by centrifugation for 30 minutes at 48,000 x *g*. The supernatant was mixed with ANTI-FLAG M2 affinity beads (Sigma-Aldrich) and incubated on a rocker for 2 hours. The resin was then washed Wash Buffer (25 mM Tris-HCl pH 8.0, 150 mM NaCl, 2 mM MgCl_2_, 2 mM ATP), then incubated with Wash Buffer supplemented with 100 μg/mL FLAG peptide (Sigma-Aldrich). The eluant was buffer-exchanged into Myosin Buffer (10 mM Tris-HCl pH 8.0, 100 mM NaCl, 2 mM MgCl_2_, and 3 mM DTT) with a 50 kDa MWCO concentrator (Amicon) via three rounds of concentrating and re-diluting with Myosin Buffer. Myosins were then flash frozen in liquid nitrogen and stored at −80°C until use.

Human profilin was purified as previously reported^6^. In brief, cells were resuspended in Lysis Buffer D (10 mM Tris-HCl pH 8.0, 1 mM EDTA, 1 mM DTT, 1 mM PMSF, 1 μg/mL aprotinin, leupeptin, and pepstatin A) and lysed with an Avestin Emulsiflex C5 homogenizer. The lysate was clarified via centrifugation for 30 minutes at 48,000 x *g*, then applied to a DEAE Sepharose Fast Flow gravity column (Cytiva). The flowthrough was collected and precipitated with 35% ammonium sulfate. The mixture was then clarified by centrifugation for 30 minutes at 48,000 x *g*. The supernatant was collected then precipitated with 61% ammonium sulfate. The precipitate was then collected via centrifugation for 30 minutes at 48,000 x *g* and resuspended in Dialysis Buffer B (5 mM potassium phosphate pH 7.5 and 1 mM DTT) and dialyzed (Spectra/Por, 3.5K MWCO) against Dialysis Buffer B overnight. The mixture was then collected, filtered through a 0.22 μm polyethersulfone (PES) membrane filter (Millipore), and applied to a 5 mL prepacked Mini ceramic hydroxyapatite (CHT) type 1 column (Bio-Rad). The flowthrough was collected and purified by SEC using a HiLoad 16/600 Superdex 200 (Cytiva) in Profilin Buffer (10 mM HEPES pH 7.0, 150 mM NaCl, and 3 mM DTT).

Actin was purified from chicken skeletal muscle as previously reported^7^. Briefly, chicken skeletal muscle acetone powder was resuspended in G-Ca buffer (2 mM Tris-Cl pH 8.0, 0.5 mM DTT, 0.2 M ATP, 0.01% NaN_3_, 0.1 mM CaCl_2_) and mixed on a nutator. Resuspended actin was ultracentrifuged in a Beckman Ti70 rotor at 42,500 RPM for 30 minutes. The supernatant was collected, and 50 mM KCl and 2 mM MgCl_2_ were added to initiate actin polymerization. After 1 hour, 0.8 M KCl was added to facilitate dissociation of actin binding factors from actin filaments. The solution was then ultracentrifuged in a Ti70 rotor at 42,500 RPM for 3 hours, and the pellet was resuspended in G-Ca buffer and incubated overnight. The mixture was homogenized in a Dounce homogenizer, then sheared through 26G and 30G needles, and finally dialyzed (SnakeSkin, 10K MWCO) in G-Ca buffer overnight. The actin mixture was sheared again through a 30G needle and then dialyzed (SnakeSkin, 10K MWCO) in fresh G-Ca buffer overnight. The protein was then ultracentrifuged in a Beckman Ti90 rotor at 70,000 RPM for 3 hours. The top 5 mL of the supernatant was collected and purified by SEC using a HiLoad 16/600 Superdex 200 column in G-Ca buffer. The second half of the peak was collected and stored at 4°C until use.

### Preparing PEG-coated coverslips

No. 1.5H 24 x 60 mm glass coverslips (Marienfeld) were sonicated in 2% Hellmanex III (Hellma) in a glass coplin staining jar for 20 minutes, followed by a MilliQ water rinse. They were then sonicated in acetone and then 1M KOH for 20 minutes each. Coverslips were rinsed with MilliQ water, then sonicated for 5 minutes in MilliQ water three times. Coverslips were then treated with Nano-Strip 2X (Cyantek) for 20 minutes, followed by thorough rinsing with MilliQ water. Eight coverslips at a time were coated with 20 mL of 1 mg/mL mPEG silane M_W_ 5K (Laysan Bio) in 96% Ethanol and 9.6 mM HCl in a 10 cm^2^ tissue culture treated dish for 16 hours while shaking at room temperature. PEG-coated coverslips were then rinsed once in 100% ethanol, and twice in MilliQ water by immersion. They were left to dry in open air for 1-3 hours before immediate use or stored at 4 °C under nitrogen in a plastic coplin staining jar. Treated coverslips were used within one week.

### Halo-tag fluorophore conjugation

Halo-ligands conjugated to Alexa Fluor 488, Janelia Fluor 549, and Janelia Fluor 646 fluorophores (Promega) were reconstituted to 1 mM in DMSO. Halo-tagged proteins were mixed with fluorophores in a 1:1 concentration ratio, then incubated on ice for 1-2 hours in the dark. After Halo-ligand conjugation reactions were complete, unreacted fluorophores were removed with micro-spin columns loaded with dye removal purification resin (Pierce) at 4 °C. Proteins were then diluted to 2-8 μM in MBOS (Motility Buffer + Oxygen Scavengers: 20 mM MOPS pH 7.4, 5 mM MgCl_2_, 0.1 mM EGTA, 50 mM KCl, 15 mM glucose, 50 mM DTT, 20 μg/mL catalase (Sigma C3515), and 100 μg/mL glucose oxidase (Sigma G6125)) + 1-4 mM ATP. Aggregates were removed by ultracentrifugation at 4 °C at 100,111 x *g* in a TLA100 rotor (Beckman). Fresh Halo-fluorophore conjugated proteins were prepared for each experiment.

### Single-filament force reconstitution assay and bundle reconstitution assay

Dual myosin motor mix was prepared by mixing 0.6 μM myosin VI and 0.1 μM myosin Va in Motility Buffer (MB: 20 mM MOPS pH 7.4, 5 mM MgCl_2_, 0.1 mM EGTA, 50 mM KCl, 3 mM DTT). Single motor myosin VI mix was composed of 0.6 μM myosin VI in MB. Single motor myosin Va mix was composed of 0.35 μM myosin Va in MB. Myosin mixes were freshly prepared for each experiment.

100% ATTO 488-labeled F-actin was prepared by polymerizing 0.9 μM ATTO 488-labeled rabbit skeletal muscle G-actin (Hypermol) in the presence of G-Mg (2 mM Tris-HCl pH 8.0, 0.5 mM DTT, 0.2 mM ATP, 0.01% NaN_3_, 0.1 mM MgCl_2_) and KMEI (50 mM KCl, 1 mM MgCl_2_, 1mM EGTA, 10 mM imidazole pH 7.0) at room temperature for 1 hour in the dark. 100% Alexa Fluor 488-labelled phallodin-F-actin was prepared by polymerizing 0.9 μM unlabeled chicken skeletal muscle G-actin in the presence of G-Mg and KMEI, followed by the addition of Alexa Fluor 488 phalloidin (Invitrogen) in a 1:1.2 actin to phalloidin molar ratio. 30% ATTO 488-labeled F-actin was prepared by co-polymerizing 0.6 μM of unlabeled chicken skeletal muscle G-actin with 0.3 μM ATTO 488-labelled rabbit skeletal muscle G-actin (Hypermol) in the presence of G-Mg and KMEI. For all experiments featuring zyxin in Figure 1, 100% ATTO-488 labeled F-actin was used. For experiments featuring FHL3, 100% Alexa Fluor 488-labelled phallodin-F-actin was used. For all other experiments, 30% ATTO 488-labeled F-actin was used. For experiments with profilin-G-actin, a working concentration of 30% ATTO 488-labeled G-actin was prepared by mixing 1.5 μM ATTO 488-labeled rabbit skeletal muscle G-actin with 3.5 μM of unlabeled chicken skeletal muscle G-actin and 20 μM of profilin in G-Mg. F-actin and G-actin mixes were freshly prepared for each experiment.

Halo-tagged repair proteins were fluorophore labelled as described above and diluted to working concentrations of 2 μM in MBOS with 1 μM calmodulin and the following ATP concentrations: experiments without profilin actin,1 mM; experiments with myosin VI and profilin actin, 2 mM; experiments with myosin Va and profilin actin, 4 mM. Proteins were clarified via ultracentrifugation at 100,111 x *g* for 12 minutes at 4 °C. After ultracentrifugation, repair protein mixes were diluted to their final concentration per experiment. Blocking buffer was prepared by mixing 1 mg/mL κ-casein (Sigma), 1 mg/mL bovine serum albumin (Gemini, BSA), and 0.1% polyvinylpyrrolidone (Sigma, PVP10, M_w_ = 10,000) in MB. All buffers, profilin-G-actin mix, and myosin mixes were centrifuged at 21,130 x *g* in a tabletop centrifuge prior to imaging.

Imaging wells were prepared by attaching a CultureWell reusable PDMS gasket (Grace Bio-Labs, 6 mm diameter and 1 mm well depth) onto a PEG-coated coverslip. Before imaging, each well was treated with the following sequence: 10 μL of myosin mix for 4 minutes, blocked with 10 μL of blocking buffer for 1 minute, 18 μL of F-actin for 30 – 60 seconds, rinsed with 18 μL of MB, and then immersed in 18 μL of MBOS for imaging via TIRF microscopy. Sequential solutions were exchanged by pipetting.

The imaging sample was mounted onto a Nikon Ti-E microscope equipped with an H-TIRF motorized module. Time-lapse multi-channel TIRF imaging was initiated to visualize F-actin in the absence of ATP (minus force), for 6 – 9 s, followed by the addition of 18 μL of repair protein mixes in MBOS + ATP. In experiments with profilin-G-actin, 18 μL repair protein mixes and 4 μL of 5 μM profilin-G-actin was added. For force reconstitution assays with zyxin alone, repair protein mix contained 50 or 500 nM zyxin alone. In VASP experiments, repair protein mix contained 50 nM zyxin and 100 nM VASP. For ɑ-actinin experiments, repair protein mix contained 50 or 500 nM zyxin and 100 nM ɑ-actinin. For bundle reconstitution assays, repair protein mix contained 50 nM zyxin, 100 nM VASP, and 100 nM ɑ-actinin in MBOS + 2 mM ATP. The final experimental concentration is half what was added to the imaging well. The final concentrations of each protein across experiments are listed in Table S1.

Images for all experiments except for those in Figure 2 were acquired at room temperature on a Nikon H-TIRF system every 3 s through a CFI Apo 100X TIRF oil immersion objective (NA 1.49), a quad filter (Chroma), and an iXON EMCCD camera (Andor) with Perfect Focus System (Nikon) engaged. In Figure 2, images were acquired every 1 s to capture VASP dynamics on zyxin patches. Illumination was provided by 488, 561, and 640 nm lasers (Agilent) switched by an acousto-optic tunable filter. Images were captured at a depth of 16-bit. Image acquisition was performed using NIS-Elements software (Nikon).

### Bundle ablation assay

Actin bundles were prepared with the bundle reconstitution protocol described above, without immediately mounting coverslips on a microscope for imaging. Coverslips were incubated for 10 minutes at room temperature, followed by 3 rinses with MB, then treated with 12 μL of 0.15 μM myosin VI in MBOS. Sequential solutions were exchanged via pipetting.

Samples were then mounted onto a Nikon Ti-2 microscope equipped with an iLas2 ring-TIRF module (Gataca System) and a micropoint laser module (Photonics Instruments). Multi-channel TIRF imaging was initiated to visualize actin bundles prior to the addition of repair proteins and ATP, followed by the addition of 28 μL of repair protein mixes in MBOS + ATP. In these experiments, repair protein mix was composed of 20 μL of 200 nM zyxin, 100 nM VASP, 200 nM ɑ-actinin in MBOS + 3 mM ATP and 8 μL of 5 μM profilin-G-actin. Once repair protein mix was added, a 1 - 5 pixel wide rectangle was drawn perpendicular to the bundle to target laser ablation.

Images were acquired every 2 seconds through a CFI Apo 100X TIRF oil immersion objective (NA 1.49), a quad filter (Chroma), and a Neo sCMOS camera (Andor) with Perfect Focus System (Nikon) engaged. Laser ablation was conducted with a 405 nm (50 mW) laser using 50-75% laser power and a 500 μs / pixel dwell time. For experiments without ɑ-actinin in the repair protein mix, 4 – 8% laser power was used. 2 – 3 frames were acquired prior to laser ablation. Illumination was provided by 488 (60 mW), 561 (50 mW), and 640 (40 mW) nm lasers. Images were captured at a depth of 16-bit using NIS-Elements software (Nikon).

### Droplet assay

For droplet experiments, Halo-tagged ɑ-actinin was labeled with AlexaFluor 488, Halo-tagged VASP was labeled with Janelia Fluor 549, and Halo-tagged zyxin was labeled with Janelia Fluor 646 as described above. Proteins were initially diluted to 4 μM in MBOS as a working concentration, then mixed at final concentrations with or without 3% PEG M_W_ 8K in MBOS in a 0.2 mL PCR tube and incubated for 10 minutes at room temperature in the dark.

Imaging wells were prepared by attaching a CultureWell reusable PDMS gasket (Grace Bio-Labs, 3 mm diameter and 1 mm well depth) onto a PEG-coated coverslip. 10 μL of protein mixture was added to wells and incubated for 20 minutes at room temperature in a dark and humid chamber to prevent evaporation. Samples were then mounted onto a Leica DMi8 microscope equipped with an instant structured illumination microscopy (VisiTech, iSIM) module and photo-bleaching/activation/ablation module (VisiTech). Single multichannel iSIM images were captured to visualize droplet distributions. For fusion and FRAP experiments, timelapse imaging was performed.

Images were acquired through a 100X oil immersion objective (NA 1.47), a quad filter (Chroma), and an Orca sCMOS camera (Hamamatsu) with hardware-based autofocus (Leica) engaged. For fusion experiments, images were acquired every 1 second. Illumination was provided by 488 (150 mW), 561 (100 mW), and 640 (100 mW)-nm lasers. Images were captured at a depth of 16-bit. Image acquisition was performed using VisiView software (Visitron Systems).

### Droplet FRAP experiments

Zyxin, VASP, and ɑ-actinin tripartite droplets were prepared at 1 μM per protein and imaged as described above. For FRAP, droplets on the surface of the glass coverslip were selected and circular regions of interest (ROI) were drawn around each target. Photobleaching was conducted with a 405 nm (100 mW) laser, with a 25 ms / pixel dwell time, illuminating each droplet twice. Images were acquired every 1 second with the same acquisition parameters as described above.

### Cell culture

A previously reported immortalized zyxin^−/−^ mouse embryonic fibroblast (Zyxin^−/−^ MEFs) cell line was generated from a tissue (torso) explant from newborn mice by the Beckerle lab^8^. The sex of the cell line is unknown because it has not been karyotyped. Zyxin^−/−^ MEFs were cultured in DMEM (Gibco, 11995073) supplemented with 4.5 g/L D-glucose and L-glutamine, 110 mg/L sodium pyruvate, 10% heat-inactivated fetal bovine serum (FBS, Sigma-Aldrich), 1X GlutaMAX (Gibco), 1X MEM non-essential amino acids (Gibco), and 0.5% antibiotic-antimycotic (Gibco) at 37 °C in 5% CO_2_.

FreeStyle 293-F cells (Gibco) were cultured in suspension with FreeStyle 293 expression medium (Gibco) on an orbital shaker at 37 °C in 8% CO_2_.

### Transient transfection

Zyxin^−/−^ MEFs were harvested at 90% confluency for transfection via nucleofection. Prior to nucleofection, 35 mm glass-bottom dishes (1.5 coverglass, Ibidi) were coated with fibronectin (EMD Millipore) for 1 hour at 37°C, then stored in culture media. 0.5 μg of F-tractin-mScarlet plasmid and 0.75 ug of Zyxin-mNeonGreen plasmid were mixed with solution SE from SE Cell Line 4D-Nucleofector S Kits (Lonza). 4×10^5^ cells were mixed with the plasmid-SE solution and nucleofected with program CM-137 using a 4D-Nucleofector (Lonza).

### Live cell Airyscan confocal imaging and laser ablation

Cells were imaged in FluoroBrite DMEM (Gibco) supplemented with 10% heat-inactivated FBS (Sigma-Aldrich), 1X MEM Non-Essential Amino Acids (Gibco), 1X GlutaMAX (Gibco), 0.5X Anti-Anti (Gibco), 20 mM Sodium Lactate (Sigma-Aldrich), and 1:100 OxyFluor (Oxyrase).

Samples were mounted onto a Zeiss Axio Observer Z1/7 microscope equipped with an Airyscan detector and FRAP module. Airyscan Confocal microscopy was performed on a Zeiss LSM 980 using a 63X oil immersion objective (NA=1.4) at 37°C and 5% CO_2_ in a stage-top incubator (PECON). Laser ablation was conducted via a 405 nm (30 mW) laser, at 20% laser power with a 500 μs / pixel dwell time. Illumination was provided by 488 nm (30 mW) and 561 nm (25 mW) lasers. Images were captured every 10 seconds at a depth of 16-bit. Image acquisition was performed using Zen Blue software (Zeiss).

## DATA ANALYSIS

Image analysis was performed with custom Python scripts utilizing functions from the scikit-image package^9^ unless otherwise specified.

### F-actin–bridge/tail interface velocity measurements

To measure velocity coupling between LIM protein bridges/tails and F-actin, contacting boundary regions in each channel were tracked with the Manual Tracker plugin in FIJI^10^. Instantaneous velocities were calculated from the distance traveled per frame versus the framerate, which were then smoothed by a rolling average of 10 frames using GraphPad Prism. Dynamics were categorized into phases based on major transitions in both the velocity and bridge growth / stability.

### Detecting and classifying F-actin-bound LIM protein patches, bridges, and tails

Analysis was adapted from our previous study of actin-bound LIM protein patches^1^. For each TIRF movie, a rolling average with a window size of 3 frames was generated for each channel to minimize intensity fluctuations when calculating masks. Averaged movies were then smoothed with a gaussian blur. A Sobel transformation was then applied to both the actin and LIM protein channels to detect the edges of actin filaments and zyxin patches. The transformed images were then thresholded (Li method for actin, Yen method for patches), followed by filling of holes. The LIM protein channel was bandpass filtered to remove objects smaller than 10 pixels and larger than 150 pixels, then each potential zyxin patch was labelled and tracked across frames. Due to non-uniform TIRF illumination and the modest signal-to-noise ratio of the zyxin channel, binding events could be lost in individual frames during tracking. Therefore, for each tracked object, both the proceeding 3 frames and following 3 frames (9 seconds) were searched for overlapping objects to account for blinking or intensity fluctuations in low signal-to-noise binding events. Persistent objects (those overlapping with at least one other labeled object in neighboring frames) were retained, then a depth first traversal was implemented to assign them as putative zyxin patches and extract their trajectories.

Putative LIM protein patches were then tracked across frames. To quantify fractional LIM protein-actin overlap, each LIM protein patch mask was multiplied by its corresponding actin mask, then the area of this overlap mask was divided by the area of the patch mask. Trajectories featuring nonzero overlap with F-actin for at least 75% of their lifetime were retained for further analysis. To account for non-uniform TIRF illumination, the local background for each patch was measured by dilating the patch mask. A per-frame normalized intensity ratio was then calculated by dividing the average LIM protein intensity within each patch by its average local background intensity. Trajectories with a maximum intensity ratio of less than 1.3 were considered too dim to be tracked and were excluded from further analysis. Detailed quantification of each patch was then performed on a per-frame basis using the skimage.measure.regionprops function. Due to the imaging inconsistencies highlighted above, some patch trajectories contained gaps or isolated outlier values, which were replaced with measurements from the frame preceding the gap / outlier.

Patches were then classified as actin-bound patches, bridges, or trails on a per-frame basis based on patterns of actin overlap. Patches featuring two detected regions of LIM protein-actin overlap were classified as bridges. Patches featuring a single region of LIM protein-actin overlap were classified as actin-bound patches if they displayed 70% fractional overlap; otherwise, they were assigned as tails. Any trajectories featuring transitions between patch classes were split based on these assignments for downstream analysis.

### Detecting and tracking interactions between actin-bound zyin patches and VASP

Analysis was conducted as described in the previous section with the following modifications. The actin channel was thresholded (Otsu method) and objects smaller than 4 pixels were removed. The zyxin mask for each frame was then multiplied by the corresponding actin mask to select for actin-bound patches. To focus on putative patches, objects were then labelled in this zyxin-actin mask and tracked across frames, and those with overlap in neighboring frames were retained. The VASP channel was thresholded (Yen method) and multiplied by the corresponding actin mask for each frame to remove non-specific VASP binding to the glass coverslip, followed by removing objects smaller than 4 pixels. To detect VASP binding events on zyxin patches, the VASP mask was multiplied by the zyxin-actin mask. To detect VASP binding events on bare F-actin, the VASP mask was multiplied by the inverse of the zyxin-actin mask. As described above, gaps in longer-lived binding event trajectories due to imaging inconsistencies were then filled by searching for overlap using a depth first traversal, followed by average intensity measurement and normalization versus local background, which is defined as “enrichment” for VASP. VASP binding events with an enrichment lower than 1 were excluded from further analysis. The maximum VASP enrichment along each binding event trajectory was then measured, followed by the average maximum VASP enrichment on bare F-actin and zyxin patches per trial.

### Quantifying lifetime of zyxin patches and zyxin-VASP binding events

The lifetime of each LIM protein patch or zyxin patch-VASP binding event was computed from the number of frames detected. For LIM protein patches, relative cumulative frequency distributions of patch lifetime were plotted to estimate half-life. For zyxin patch-VASP binding events, two categories were observed: (1) transient, with a lifetime of 1 – 2 seconds (as previously^11^), and (2) stable, with a lifetime of tens of seconds to minutes. Because transient binding events predominated, and our temporal resolution (1 frame / second) was similar to their lifetime, the half-life of all zyxin patch-VASP binding events could not be accurately estimated. Therefore, we compared the average lifetimes observed across experimental conditions, which captures lifetime variations in long-lived binding events.

### Detecting and tracking ɑ-actinin on zyxin patches in the presence of myosin forces (+ATP)

Analysis was performed similarly as for zyxin patch-VASP binding interactions, with the following modifications. The actin channel was thresholded by the Yen method and objects smaller than 10 pixels were removed. The zyxin channel was thresholded by the triangle method, and the ɑ-actinin channel was threshold by the Yen method. Both were multiplied by the actin mask, and objects smaller than 4 pixels were removed. Tracking and quantification was performed as described above, except a window of 15 seconds was searched, and ɑ-actinin binding events with an enrichment below 1.3 were excluded from further analysis.

### Detecting and quantifying ɑ-actinin-zyxin interactions in the absence of myosin forces (-ATP)

The last frame of each 5-minute movie was used for quantification, as we observed that zyxin stably associated with non-dynamic ɑ-actinin clusters on F-actin. The actin channel was thresholded (Li method), and objects smaller than 5 pixels were removed. Due to the low signal-to-noise ratio of zyxin binding events, Sobel transformation was applied to the zyxin channel to detect cluster edges. The transformed images were then thresholded (Yen method), followed by hole filling. Because Sobel transformation smooths edges, the resultant masks were larger than the actual size of clusters, and they were therefore eroded once. The actin mask was then applied to the zyxin mask, and objects smaller than 3 pixels were removed. The ɑ-actinin channel was also sequentially Sobel transformed, thresholded (Li method for ɑ-actinin^WT^; Yen method for ɑ-actinin^ABM^), filled, then eroded. The actin mask was then applied to the ɑ-actinin mask and objects smaller than 3 pixels were removed. To isolate zyxin-ɑ-actinin binding events, the zyxin mask was multiplied by the ɑ-actinin mask. ɑ-actinin enrichment was quantified as described above.

### Actin fold increase at VASP-enriched zyxin patches

A 20 x 20 pixel box was drawn around each VASP-enriched zyxin patch, and all channels were subjected to rolling ball background subtraction in FIJI. Equal numbers of control F-actin regions lacking zyxin patches were also selected. The per-frame integrated density of the actin channel (sum of intensity across all pixels in the box) was then quantified. Actin fold change was calculated by dividing the integrated density of each frame versus the average integrated density of the first 5 frames. The time-averaged actin fold change was calculated for each region over the imaging period, and paired analysis was conducted between zyxin patches and control regions from the same field of view (Figure 4D). The five zyxin patch and control F-actin regions with the maximum average actin fold change were selected for plotting in Figure 4C, displaying a rolling average of 10 frames (30 s).

### Droplet size quantification

For each image, all three channels (zyxin, VASP, and ɑ-actinin) were thresholded (Otsu method) and objects smaller than 20 pixels were removed. All three masks were multiplied together to generate a droplet mask, from which the droplet area was computed. The fraction plotted in Figure S5C was computed by dividing the droplet area by the total area of the image.

### FRAP analysis

All FRAP analysis was conducted with the Stowers Plugins in FIJI. Full field movies were bleach corrected by histogram matching^12^ to account for bulk photobleaching during imaging. As each field of view was 1440 x 965 pixels (94 x 62 microns) featuring dozens of condensates, we found that targeted photobleaching of 6-8 condensates (less than 10% of all droplets) did not detrimentally impact the histogram matching algorithm. Bleached regions of interest (ROI) were selected using a circular selection tool, and average ROI intensity was measured and plotted against time with the plugin ‘create spectrum jru v1.’ Traces of individual FRAP ROIs were grouped with ‘combine trajectories jru v1’ and normalized to the minimum and maximum of each trace with ‘normalize trajectories jru v1.’ Normalized traces were plotted using GraphPad Prism.

### Quantifying protein enrichment in droplets

For zyxin / VASP / α-actinin tripartite droplets and zyxin / VASP bi-partite droplets, all channels were thresholded (Otsu method) to generate masks, and objects smaller than 20 pixels were removed. For zyxin / ɑ-actinin bipartite droplets, rolling ball background subtracted images were used to generate masks for both channels, and each mask was then applied to its corresponding raw channel. For experiments featuring zyxin^LDO^, the VASP or ɑ-actinin mask was applied to the zyxin channel to quantify zyxin^LDO^ enrichment, as the zyxin^LDO^ enrichment was too low to generate well-defined masks. To quantify zyxin enrichment within droplets, the average intensity inside each masked object was divided by the average intensity of its local background, calculated by dilating the mask. Droplets smaller than 40 x 40 pixels (0.18 μm^2^) or featuring zyxin enrichment lower than 1.1 were excluded from further analysis.

### Quantification of bundle thickness

To quantify bundle thickness over time, the actin channel of each movie was thresholded (Otsu method), and objects smaller than 50 pixels were removed. The bundle thickness index of each frame (adapted from a published approach^13^) was calculated as the number of binary erosions necessary to reduce the area of the actin mask of that frame to the same area or less than that of the first frame.

### Quantification of reconstituted strain sites

To quantify repair frequency, individual reconstituted strain sites were manually divided into two categories: 1) events that broke or did not repair, or 2) events which repaired. A subset of strain sites which remained in a stable position during their lifetime were selected, and line scans were performed across their zyxin flashes to generate kymographs.

To quantify actin recovery, the average actin intensity of the kymographs at each time point was additionally quantified and normalized to the maximum and minimum intensity. To quantify VASP fold increase, the average VASP intensity of the kymographs at each timepoint was additionally quantified and normalized to the average VASP intensity pre-ablation. Strain sites that exhibited significant drift were excluded from fluorescence intensity over time plots due to poor kymograph quality, but they were still included in repair frequency quantification.

### Quantifying actin recovery rate at stress fiber strain sites (SFSSs)

Movies were bleach corrected via histogram matching with FIJI. SFSSs were then cropped out of the field of view and registered by rigid body transformation with the HyperStackReg plugin in FIJI^14^.

SFSSs that successfully repaired were manually divided into four actin repair path categories: recovery 1) evenly across the zyxin flash, 2) primarily from the center, (3) from one end and the center, and (4) from both ends of the zyxin flash. SFSS that primarily recovered from the center and maintained a stable length over the repair period were selected for detailed analysis. Line scans were performed across each zyxin flash to generate kymographs with FIJI. The length of the strain site was then measured. The left end and right end were designated as the first and last 10% of the SFSS, respectively, while the center of the SFSS was designated as the middle 80%. Line scans of the kymograph along the time axis with widths corresponding to each respective segment (left end, right end, center) were performed, and the average fluorescence of each segment was plotted over time. Simple linear regression was performed to estimate the recovery rate, and paired comparisons were performed between the average of the recovery rates of the two ends and the center for each SFSS.

### Statistical Analysis

All statistical analysis and plotting were performed in GraphPad Prism. Figure legends contain details of the statistical analyses throughout the study.

